# Reporting animal research: Explanation and Elaboration for the ARRIVE guidelines 2019

**DOI:** 10.1101/703355

**Authors:** Nathalie Percie du Sert, Amrita Ahluwalia, Sabina Alam, Marc T. Avey, Monya Baker, William J. Browne, Alejandra Clark, Innes C. Cuthill, Ulrich Dirnagl, Michael Emerson, Paul Garner, Stephen T. Holgate, David W. Howells, Viki Hurst, Natasha A. Karp, Katie Lidster, Catriona J. MacCallum, Malcolm Macleod, Esther J Pearl, Ole Petersen, Frances Rawle, Penny Reynolds, Kieron Rooney, Emily S. Sena, Shai D. Silberberg, Thomas Steckler, Hanno Würbel

## Abstract

Improving the reproducibility of biomedical research is a major challenge. Transparent and accurate reporting are vital to this process; it allows readers to assess the reliability of the findings, and repeat or build upon the work of other researchers. The NC3Rs developed the ARRIVE guidelines in 2010 to help authors and journals identify the minimum information necessary to report in publications describing *in vivo* experiments.

Despite widespread endorsement by the scientific community, the impact of the ARRIVE guidelines on the transparency of reporting in animal research publications has been limited. We have revised the ARRIVE guidelines to update them and facilitate their use in practice. The revised guidelines are published alongside this paper. This Explanation and Elaboration document was developed as part of the revision. It provides further information about each of the 21 items in ARRIVE 2019, including the rationale and supporting evidence for their inclusion in the guidelines, elaboration of details to report, and examples of good reporting from the published literature.

## Introduction

Transparent and accurate reporting is essential to improve the reproducibility of scientific research; it enables others to scrutinise the methodological rigour of the studies, assess how reliable the findings are, and repeat or build upon the work.

However, evidence shows that the majority of publications fail to include key information and there is significant scope to improve the reporting of studies involving animal research [1–4]. To that end, the NC3Rs published the ARRIVE guidelines in 2010. The guidelines are a checklist of information to include in a manuscript to ensure that publications contain enough information to add to the knowledge base [5]. The guidelines have received widespread endorsement from the scientific community and are currently recommended by more than a thousand journals, with further endorsement from research funders, universities and learned societies worldwide.

Studies measuring the impact of ARRIVE on the quality of reporting have produced mixed results [6–11] and there is evidence that *in vivo* scientists are not sufficiently aware of the importance of reporting the information covered in the guidelines, and fail to appreciate the relevance to their work or their research field [12].

As a new international working group – the authors of this publication, we have revised the guidelines to update them and facilitate their uptake; the ARRIVE guidelines 2019 are published alongside this paper [13]. We have updated the recommendations in line with current best practice, reorganised the information and classified the items into two sets. The ARRIVE Essential 10 constitute the minimum reporting requirement and the Recommended Set provides further context to the study described. The two sets help authors, journal staff, editors and reviewers use the guidelines in practice, and allow a pragmatic implementation with an initial focus on the most critical issues. Once the Essential 10 are consistently reported in manuscripts, items from the Recommended Set can be added to journal requirements over time until all 21 items are routinely reported in all manuscripts. Full methodology for the revision and the allocation of items into sets is described in the accompanying publication [13].

A key aspect of the revision was to develop this Explanation and Elaboration document to provide background and rationale for each of the 21 items of ARRIVE 2019. Here we present additional guidance for each item and subitem, explain the importance of reporting this information in manuscripts that describe animal research, elaborate on what to report, and provide supporting evidence. Each subitem is also illustrated with examples of good reporting from the published literature.

### Box 1: Glossary

**Bias:** Introduction of a systematic error in the estimated effect of an intervention, caused by inadequacies in the design, conduct, or analysis of an experiment.

**Effect size:** Quantitative measure that estimates the magnitude of differences between groups, or relationships between variables.

**Experimental unit:** Biological entity subjected to an intervention independently of all other units, such that it is possible to assign any two experimental units to different treatment groups.

**External validity:** Extent to which the results of an animal experiment provide a correct basis for generalisations to other populations of animals (including humans) and/or other environmental conditions.

**False positive:** Statistically significant result obtained by chance when the effect being investigated does not exist.

**False negative:** Non-statistically significant result obtained when the effect being investigated genuinely exists.

**Independent variable of interest:** Factor that a researcher manipulates within a controlled environment in order to test its impact on the outcome measured. Also known as: predictor variable, factor of interest.

**Internal validity:** Refers to the rigour of the study design and statistical analysis to isolate cause and effect, and attribute the effect observed to manipulation of the independent variable of interest. In an experiment with high internal validity, sources of bias and chance observations are minimised. In an experiment with low internal validity, the effect may be caused by bias, chance and other nuisance variables rather than the independent variable(s) of interest.

**Null and alternative hypotheses:** The null hypothesis (H_0_) refers to the postulate that the response being measured is unaffected by the experimental manipulation being tested. The alternative hypothesis (H_1_) refers to the postulate that manipulating the independent variable of interest has an effect on the response measured.

**Nuisance variable:** Sources of variability or conditions that could potentially bias results. Also known as: confounding factor, confounding variable

**Outcome measure:** Any variable recorded during a study to assess the effects of a treatment or experimental intervention. Also known as: dependent variable, response variable

**Power:** Probability that a test of significance will detect an effect (i.e. a deviation from the null hypothesis), if an effect exists (i.e. true positive result).

**Sample size:** Number of experimental units per group, also referred to as N number.

Definitions adapted from [14, 15] and placed in the context of animal research.

## 1. ARRIVE Essential 10

The ARRIVE Essential 10 (Box 2) constitute the minimum reporting requirement, to ensure that reviewers and readers can assess the reliability of the findings presented. There is no ranking within the set, items are presented in a logical order.

### Box 2: ARRIVE Essential 10

1. Study design
2. Sample size
3. Inclusion and exclusion criteria
4. Randomisation
5. Blinding
6. Outcome measures
7. Statistical methods
8. Experimental animals
9. Experimental procedures
10. Results

### Item 1. Study design

For each experiment, provide brief details of study design including:

1a. The groups being compared, including control groups. If no control group has been used, the rationale should be stated.

#### Explanation

The choice of control or comparator group is dependent on the experimental objective. Negative controls are used to determine if a difference between groups is caused by the intervention (e.g. wild-type animals vs genetically modified animals, placebo vs active treatment, sham surgery vs surgical intervention). Positive controls can be used to support the interpretation of negative results or determine if an expected effect is detectable.

It may not be necessary to include a separate control with no active treatment if, for example, the experiment aims to compare a treatment administered by different methods (e.g. intraperitoneal administration vs. oral gavage), or animals that are used as their own control in a longitudinal study. A pilot study, such as one designed to test the feasibility of a procedure might also not require a control group.

For complex study designs, a visual representation is more easily interpreted than a text description, so a timeline diagram or flow chart is recommended. Diagrams facilitate the identification of which treatments and procedures were applied to specific animals or groups of animals, and at what point in the study these were performed. They also help to communicate complex design features such as clustering or nesting (hierarchical designs), blocking (to reduce unwanted variation, see **item 4 – Randomisation**), or repeated measurements over time on the same experimental unit (repeated measures designs) [16, 17]. The Experimental Design Assistant (EDA) is a platform to support researchers in the design of in vivo experiments, it can be used to generate diagrams to represent any type of experimental design [18].

Report the groups clearly so that test groups, comparators and controls (negative or positive) can be identified easily. State clearly if the same control group was used for multiple experiments.

#### Examples

1. *“The DAV1 study is a one-way, two-period crossover trial with 16 piglets receiving amoxicillin and placebo at period 1 and only amoxicillin at period 2. Amoxicillin was administered orally with a single dose of 30 mg.kg^−1^. Plasma amoxicillin concentrations were collected at same sampling times at each period: 0.5, 1, 1.5, 2, 4, 6, 8, 10 and 12 h.” [19]*
2. *“Figure A: Example of a study plan created using the Experimental Design Assistant showing a simple comparative study for the effect of two drugs on the metastatic spread of two different cancer cell lines. Block randomisation has been used to create 3 groups containing an equal number of zebrafish embryos injected with either cell line, and each group will be treated with a different drug treatment (including vehicle control). Each measurement outcome will be analysed by 2-way ANOVA to determine the effect of drug treatment on growth, survival and invasion of each cancer cell line.” [20]*

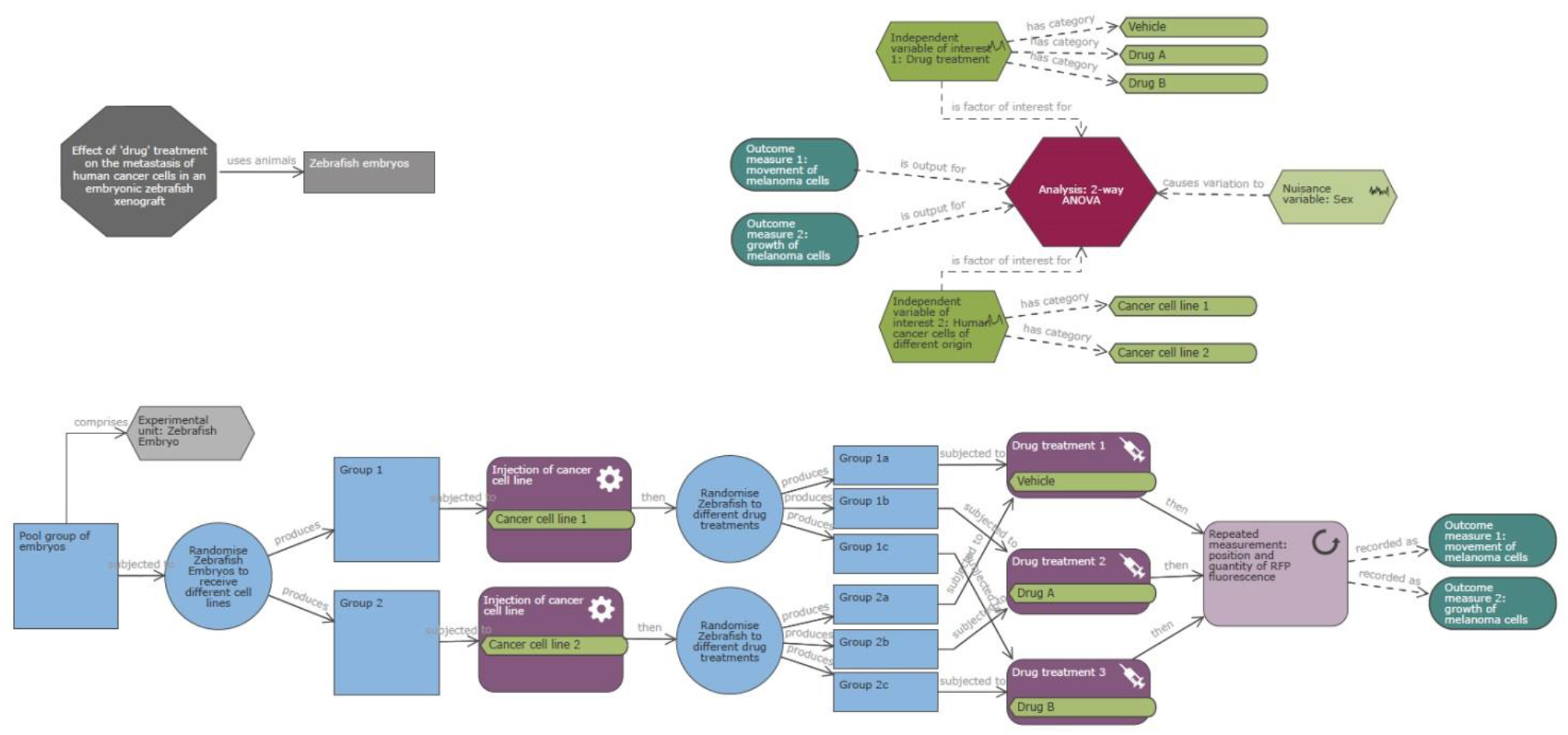

1b. The experimental unit (e.g. a single animal, litter, or cage of animals).

#### Explanation

The experimental unit is the biological entity subjected to an intervention independently of all other units, such that it is possible to assign any two experimental units to different treatment groups. The sample size is the number of experimental units per group.

Clearly indicate the experimental unit for each experiment so that the sample sizes and statistical analyses can be properly evaluated. There is a risk that if the experimental unit is not correctly identified, the sample size used in the data analysis will be incorrect. Inflation of the sample size by conflating experimental units with subsamples or repeated measurements is known as ‘pseudoreplication’. This may invalidate the analysis and resulting conclusions [21, 22] (see also **item 7 – Statistical methods**).

Commonly, the experimental unit is the individual animal, each independently allocated to a treatment group (e.g. a drug administered by injection). However, the experimental unit may be the cage or the litter (e.g. a diet administered to a whole cage, or a treatment administered to a dam and investigated in her pups), or it could be part of the animal (e.g. different drug treatments applied topically to distinct body regions of the same animal). Animals may also serve as their own controls receiving different treatments separated by washout periods; here the experimental unit is an animal for a period of time. There may also be multiple experimental units in a single experiment, such as when a treatment such as diet is given to a pregnant dam and then the weaned pups are allocated to different diets [23]. See [24–26] for further guidance on identifying experimental units.

#### Examples

1. *“The present study used the tissues collected at E15.5 from dams fed the 1X choline and 4X choline diets (n = 3 dams per group, per fetal sex; total n = 12 dams). To ensure statistical independence, only one placenta (either male or female) from each dam was used for each experiment. Each placenta, therefore, was considered to be an experimental unit.” [27]*
2. *“We have used data collected from high-throughput phenotyping, which is based on a pipeline concept where a mouse is characterized by a series of standardized and validated tests underpinned by standard operating procedures (SOPs)…. The individual mouse was considered the experimental unit within the studies.” [28]*
3. *“Fish were divided in two groups according to weight (0.7-1.2 g and 1.3-1.7 g) and randomly stocked (at a density of 15 fish per experimental unit) in 24 plastic tanks holding 60 L of water.” [29]*
4. *“In the study, n refers to number of animals, with five acquisitions from each [corticostriatal] slice, with a maximum of three slices obtained from each experimental animal used for each protocol (six animals each group).” [30]*

### Item 2. Sample size

2a. Specify the exact number of experimental units allocated to each group, and the total number in each experiment. Also indicate the total number of animals used.

#### Explanation

Sample size relates to the number of experimental units in each group at the start of the study, and is usually represented by N (See **item 1 - Study design** for further guidance on identifying and reporting experimental units). This information is crucial to assess the validity of the statistical model and the robustness of the experimental results.

Report the exact value of N per group and the total number in each experiment. If the experimental unit is not the animal, also report the total number of animals to help readers understand the study design. For example, in a study investigating diet using cages of animals housed in pairs, the number of animals is double the number of experimental units. Reporting the total number of animals is also useful to identify if any were re-used between experiments.

#### Example

1. *“Dams (for n see Table …) were assigned to treatments in a manner that provided similar means and variances in body weight before dosing was initiated…. For statistical purposes, the numbers/group are the number of litters, not the number of pups.” [31]*

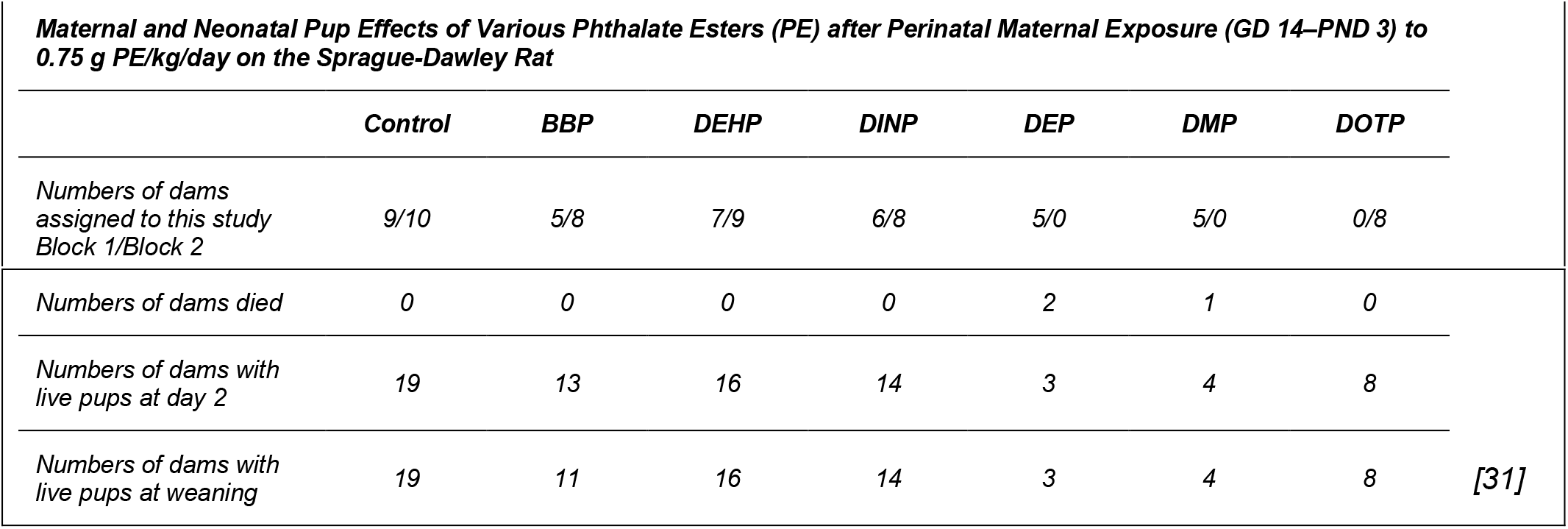

|  | Control | BBP | DEHP | DINP | DEP | DMP | DOTP |
| --- | --- | --- | --- | --- | --- | --- | --- |
| Numbers of dams assigned to this study<br>Block 1/Block 2 | 9/10 | 5/8 | 7/9 | 6/8 | 5/0 | 5/0 | 0/8 |
| Numbers of dams died | 0 | 0 | 0 | 0 | 2 | 1 | 0 |
| Numbers of dams with live pups at day 2 | 19 | 13 | 16 | 14 | 3 | 4 | 8 |
| Numbers of dams with live pups at weaning | 19 | 11 | 16 | 14 | 3 | 4 | 8 |

2b. Explain how the sample size was decided. Provide details of any *a priori* sample size calculation, if done.

#### Explanation

For any type of experiment, it is crucial to explain how the sample size was determined. For hypothesis-testing experiments, where inferential statistics are used to estimate the size of the effect and to determine the weight of evidence against the null hypothesis, the sample size needs to be justified to ensure experiments are of an optimal size to test the research question [32, 33] (see **item 13 – Objectives**). Power is the probability that a test of significance will detect an effect (i.e. a deviation from the null hypothesis), when the effect being investigated genuinely exists (i.e. true positive result). Sample sizes that are too small (i.e. underpowered studies) produce inconclusive results, whereas sample sizes that are too large (i.e. overpowered studies) raise ethical issues over unnecessary use of animals and may produce trivial findings that are statistically significant but not biologically relevant [34]. Low power has three effects: first, within the experiment, real effects are more likely to be missed; second, where an effect is detected, this will often be an over-estimation of the true effect size [25]; and finally, when low power is combined with publication bias, there is an increase in the false positive rate in the published literature [35]. Consequently, low powered studies contribute to the poor internal validity of research and risk wasting animals used in inconclusive research [36].

Study design can influence the statistical power of an experiment. Split-plot designs [37], factorial designs [38], or group-sequential designs [39] can increase the power of a study for a given number of animals. Statistical programs to help perform *a priori* sample size calculations exist for a variety of experimental designs and statistical analyses, for example G*power [40]. Choosing the appropriate calculator or algorithm to use depends on the type of outcome measures and independent variables, and the number of groups. Consultation with a statistician is recommended, especially when the experimental design is complex or unusual.

Where the experiment tests the effect of an intervention on the mean of a continuous outcome measure, the sample size can be calculated *a priori*, based on a mathematical relationship between the desired effect size, variability estimated from prior data, chosen significance level, power and sample size (See Box 3, and [24, 41] for practical advice). For an *a priori* sample size determination, report the analysis method (e.g. two-tailed student’s t-test with a 0.05 significance threshold), the effect size of interest and a justification explaining why this effect size is relevant, the estimate of variability used (e.g. standard deviation) and how it was estimated, and the power selected.

##### Box 3: Information used in a power calculation

Sample size calculation is based on a mathematical relationship between the following parameters: effect size, variability, significance level, power and sample size. Questions to consider are:

**<u>The primary objective of the experiment – what is the main outcome measure?</u>**

The primary outcome measure should be identified in the planning stage of the experiment; it is the outcome of greatest importance, which will answer the main experimental question.

**<u>What is a biologically or clinically relevant effect size?</u>**

The effect size is the minimum change in the primary outcome measure between the groups under study, which would be of interest biologically and would be worth taking forward into further work.

**<u>What is the estimate of variability?</u>**

Estimates of variability can be obtained:

- From data collected from a preliminary experiment conducted under identical conditions to the planned experiment, e.g. a previous experiment in the same lab, testing the same treatment, under similar conditions, on animals with the same characteristics.
- From the control group in a previous experiment testing a different treatment.
- From a similar experiment reported in the literature.

**<u>What risk of a false positive is acceptable? (significance threshold)</u>**

The significance level or threshold (α) is the probability of obtaining a significant result by chance (a false positive) when the null hypothesis is true (i.e. there is no real, biologically relevant difference between the groups). If it is set at 0.05 then the risk of obtaining a false positive is 1 in 20 for a single statistical test. However, the threshold or the p values will need to be adjusted in scenarios of multiple testing (e.g. by using a Bonferroni correction).

**<u>What risk of a false negative is acceptable? (power)</u>**

The power (1-β) is the probability that the experiment will correctly lead to the rejection of the null hypothesis if the effect being investigated genuinely exists (i.e. detect that there is a biologically meaningful difference when there is one). A target power between 80-95% is normally deemed acceptable.

**<u>Will you use a one or two-sided test? (directionality)</u>**

The directionality of a test depends on the distribution of the test statistics for a given analysis. For tests based on t or z distributions (such as t-tests), whether the data will be analysed using a one or two-sided test relates to whether the alternative hypothesis (H_1_) is directional or not. An experiment with a directional (one-sided) H_1_ can be powered and analysed with a one-sided test. This assumes that direction of the effect is known (this is very rare in biology) and the goal is to maximise the chances of detecting this effect. However, the investigator cannot then test for the possibility of missing an effect in the untested direction. Choosing a one-tailed test for the sole purpose of attaining statistical significance is not appropriate.

Two-sided tests with a non-directional H_1_ are much more common and allow researchers to detect the effect of a treatment regardless of its direction.

Note that analyses such as ANOVA and chi-square are based on asymmetrical distributions (F-distribution and chi-square distribution) with only one tail. Therefore, these tests do not have a directionality option.

There are several types of studies where *a priori* sample size calculations are not appropriate. For example, the number of animals needed for antibody or tissue production is determined by the amount required and the production ability of an individual animal. For studies where the outcome is a successful generation of a sample or a condition (e.g. the production of transgenic animals), the number of animals is determined by the probability of success of the experimental procedure.

In early feasibility or pilot studies, the number of animals required depends on the purpose of the study. Where the objective of the preliminary study is to improve procedures and equipment, the number of animals needed is generally small. In such cases power calculations are not appropriate and sample sizes can be estimated based on operational capacity and constraints [42]. Pilot studies alone are unlikely to provide adequate data on variability for a power calculation for future experiments. Systematic reviews and previous studies are more appropriate sources of information on variability [43].

Regardless of whether a power calculation was used or not, when explaining how the sample size was determined take into consideration any anticipated loss of animals or data, for example due to exclusion criteria established upfront or expected attrition (see **item 3 – inclusion and exclusion criteria**).

#### Examples

1. *“The sample size calculation was based on postoperative pain numerical rating scale (NRS) scores after administration of buprenorphine (NRS AUC mean = 2.70; noninferiority limit = 0.54; standard deviation = 0.66) as the reference treatment and also Glasgow Composite Pain Scale (GCPS) scores using online software (Experimental design assistant; https://eda.nc3rs.org.uk/eda/login/auth). The power of the experiment was set to 80%. A total of 20 dogs per group were considered necessary.” [44]*
2. *“We selected a small sample size because the bioglass prototype was evaluated in vivo for the first time in the present study, and therefore, the initial intention was to gather basic evidence regarding the use of this biomaterial in more complex experimental designs.” [45]*

### Item 3. Inclusion and exclusion criteria

3a. Describe any criteria established *a priori* for including or excluding animals (or experimental units) during the experiment, and data points during the analysis.

#### Explanation

Inclusion and exclusion criteria define the eligibility or disqualification of animals and data once the study has commenced. To ensure scientific rigour, the criteria should be defined before the experiment starts and data are collected [46]. Inclusion criteria should not be confused with animal characteristics (see **item 8 – Experimental animals**) but can be related to these (e.g. body weights must be within a certain range for a particular procedure) or related to other study parameters (e.g. task performance has to exceed a given threshold). Exclusion criteria may result from technical or welfare issues such as complications anticipated during surgery, or circumstances where test procedures might be compromised (e.g. development of motor impairments that could affect behavioural measurements). Criteria for excluding samples or data include failure to meet quality control standards, such as insufficient sample volumes, unacceptable levels of contaminants, poor histological quality, etc. Similarly, how the researcher will define and handle data outliers during the analysis should be also decided before the experiment starts (see subitem 3b for guidance on responsible data cleaning).

Exclusion criteria may also reflect the ethical principles of a study in line with its humane endpoints (see **item 16 – Animal care and monitoring**). For example, in cancer studies an animal might be dropped from the study and euthanised before the predetermined time point if the size of a subcutaneous tumour exceeds a specific volume [47]. If losses are anticipated, these should be considered when determining the number of animals to include in the study (see **item 2 – Sample size**).

Best practice is to include all *a priori* inclusion and exclusion/outlier criteria in a pre-registered protocol (see **item 19 – Protocol registration**). At the very least these criteria should be documented in a lab notebook and reported in manuscripts, explicitly stating that the criteria were defined before any data was collected.

#### Example

1. *“The animals were included in the study if they underwent successful MCA occlusion (MCAo), defined by a 60% or greater drop in cerebral blood flow seen with laser Doppler flowmetry. The animals were excluded if insertion of the thread resulted in perforation of the vessel wall (determined by the presence of sub-arachnoid blood at the time of sacrifice), if the silicon tip of the thread became dislodged during withdrawal, or if the animal died prematurely, preventing the collection of behavioral and histological data.” [48]*

3b. For each experimental group, report any animals, experimental units or data points not included in the analysis and explain why.

#### Explanation

Animals, experimental units, or data points that are unaccounted for can lead to instances where conclusions cannot be supported by the raw data [49]. Reporting exclusions and attritions provides valuable information to other investigators evaluating the results, or who intend to repeat the experiment or test the intervention in other species. It may also provide important safety information for human trials (e.g. exclusions related to adverse effects).

There are many legitimate reasons for experimental attrition, some of which are anticipated and controlled for in advance (see subitem 3a on defining exclusion and inclusion criteria) but some data loss might not be anticipated. For example, data points may be excluded from analyses due to an animal receiving the wrong treatment, unexpected drug toxicity, infections or diseases unrelated to the experiment, sampling errors (e.g. a malfunctioning assay that produced a spurious result, inadequate calibration of equipment), or other human error (e.g. forgetting to switch on equipment for a recording).

In some instances, it may be scientifically justifiable to remove outlier data points from an analysis, such as readings that are outside a plausible range. Providing the reasoning for removing data points enables the distinction to be made between responsible data cleaning and data manipulation. When reasons are not disclosed the reliability of the conclusions is in question, as inappropriate data cleaning has the potential to bias study outcomes [50].

There is a movement towards greater data sharing (see **item 20 – Data access**), along with an increase in strategies such as code sharing to enable analysis replication. These practices, however transparent, still need to be accompanied by a disclosure on the reasoning for data cleaning, and whether it was pre-defined.

Report all animal exclusions and loss of data points, along with the rationale for their exclusion. Accompanying these criteria should be an explicit description of whether researchers were blinded to the group allocations when data or animals were excluded (see **item 5 – Blinding** and [51]), and whether these criteria were decided prior to the experiment [8, 32, 52]. Explicitly state where built-in models in statistics packages have been used to remove outliers (e.g. GraphPad Prism’s outlier test).

#### Examples

1. *“Pen was the experimental unit for all data. One entire pen (ZnAA90) was removed as an outlier from both Pre-RAC and RAC periods for poor performance caused by illness unrelated to treatment…. Outliers were determined using Cook’s D statistic and removed if Cook’s D > 0.5. One steer was determined to be an outlier for day 48 liver biopsy TM and data were removed.” [53]*
2. *“Seventy-two SHRs were randomized into the study, of which 13 did not meet our inclusion and exclusion criteria because the drop in cerebral blood flow at occlusion did not reach 60% (seven animals), postoperative death (one animal: autopsy unable to identify the cause of death), haemorrhage during thread insertion (one animal), and disconnection of the silicon tip of the thread during withdrawal, making the permanence of reperfusion uncertain (four animals). A total of 59 animals were therefore included in the analysis of infarct volume in this study. In error, three animals were sacrificed before their final assessment of neurobehavioral score: one from the normothermia/water group and two from the hypothermia/pethidine group. These errors occurred blinded to treatment group allocation. A total of 56 animals were therefore included in the analysis of neurobehavioral score.” [48]*
3. *“Fig 1. Flow chart showing the experimental protocol with the number of animals used, died and included in the study…. After baseline CMR and echocardiography, MI was induced by left anterior descending (LAD) coronary artery ligation (n = 48), as previously described. As control of surgery procedure, sham operated mice underwent thoracotomy and pericardiotomy without coronary artery ligation (n = 12).”[54]*

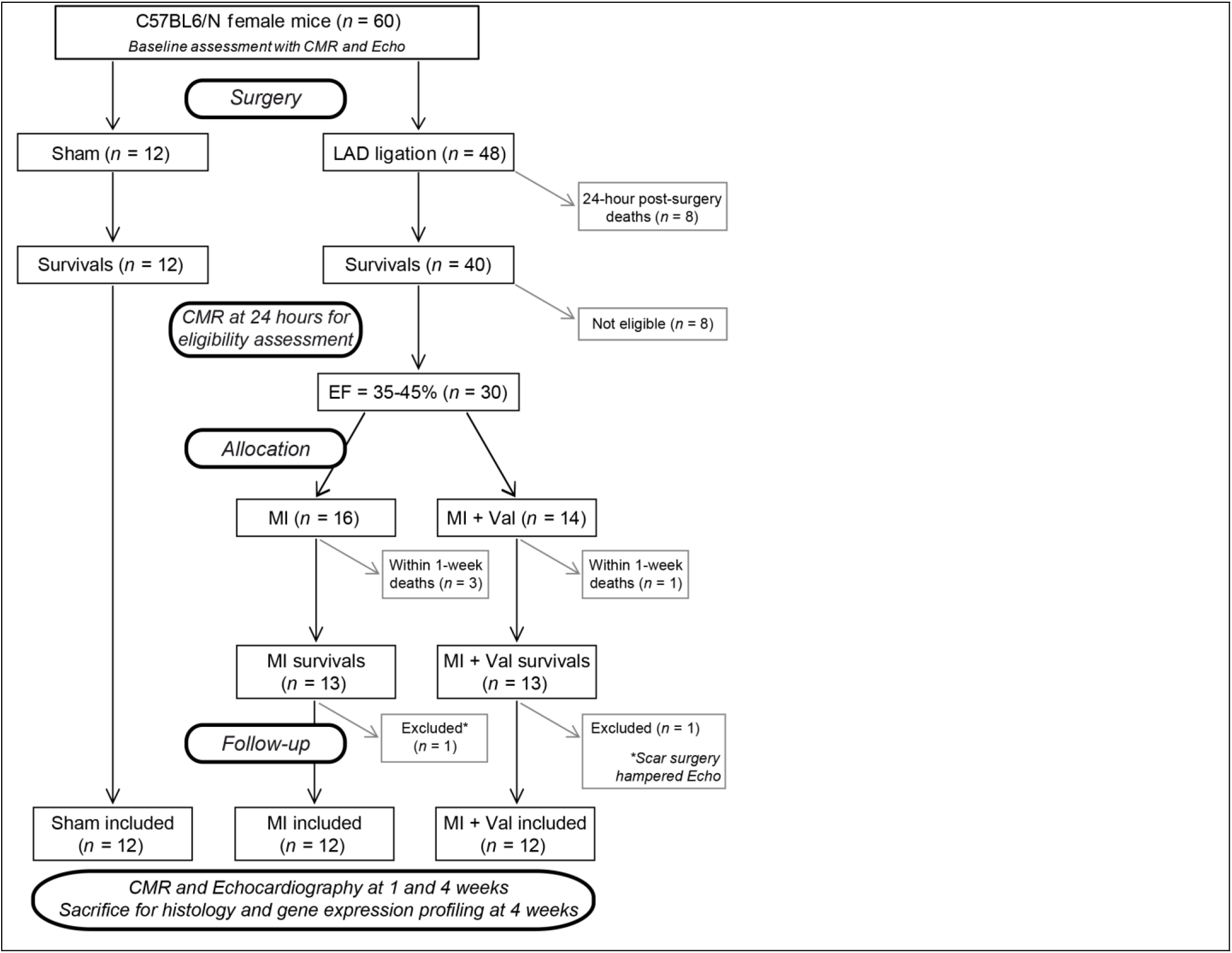

3c. For each analysis, report the exact value of N in each experimental group.

#### Explanation

The exact number of experimental units analysed in each group (i.e. the N number) is essential information for the reader to interpret the analysis, it should be reported unambiguously. All animals and data used in the experiment should be accounted for in the data presented. Sometimes, for good reasons, animals may need to be excluded from a study (e.g. illness or mortality), and data points excluded from analyses (e.g. biologically implausible values). Reporting losses will help the reader to understand the experimental design process, replicate methods, and provide adequate tracking of animal numbers in a study, especially when sample size numbers in the analyses do not match the original group numbers.

Indicate numbers clearly within the text or on figures, and provide absolute numbers (e.g. 10/20, not 50%). For studies where animals are measured at different time points, explicitly report the full description of which animals undergo measurement, and when [32].

#### Examples

1. *“Group F contained 29 adult males and 58 adult females in 2010 (n = 87), and 32 adult males and 66 adult females in 2011 (n = 98). The increase in female numbers was due to maturation of juveniles to adults. Females belonged to three matrilines, and there were no major shifts in rank in the male hierarchy. Six mid to low ranking individuals died and were excluded from analyses, as were five mid-ranking males who emigrated from the group at the beginning of 2011.” [55]*
2. *“The proportion of test time that animals spent interacting with the handler (sniffed the gloved hand or tunnel, made paw contact, climbed on, or entered the handling tunnel) was measured from DVD recordings. This was then averaged across the two mice in each cage as they were tested together and their behaviour was not independent…. Mice handled with the home cage tunnel spent a much greater proportion of the test interacting with the handler (mean ± s.e.m., 39.8 ± 5.2 percent time of 60 s test, n = 8 cages) than those handled by tail (6.4 ± 2.0 percent time, n = 8 cages), while those handled by cupping showed intermediate levels of voluntary interaction (27.6 ± 7.1 percent time, n = 8 cages).” [56]*

### Item 4. Randomisation

Describe the methods used:

4a. To allocate experimental units to control and treatment groups. If randomisation was used, provide the method of randomisation.

#### Explanation

Using randomisation during the allocation to groups ensures that each experimental unit has an equal probability of receiving a particular treatment. It helps minimise selection bias and reduce systematic differences in the characteristics of animals allocated to different groups [57–59]. However, investigators frequently confuse “random” with “haphazard” or “arbitrary” allocation. Non-random allocation can introduce bias that influences the results, as a researcher may (consciously or subconsciously) make judgements in allocating an animal to a particular group, or because of unknown and uncontrolled differences in the experimental conditions or animals in different groups. Systematic reviews have shown that animal experiments that do not report randomisation or other bias-reducing measures such as blinding, are more likely to report exaggerated effects that meet conventional measures of statistical significance [60, 61]. It is especially important to use methods of randomisation in situations where it is not possible to blind all or parts of the experiment but even with randomisation, researcher bias can pervert the allocation. This can be avoided by using allocation concealment (**see item 5 – Blinding**). In studies where sample sizes are small, simple randomisation may result in unbalanced groups; here randomisation strategies to balance groups such as randomising in matched pairs [62–64] and blocking are encouraged [24].

Report the type of randomisation used (simple, stratified, randomised complete blocks, etc.; see Box 4), the method of randomisation (e.g. computer-generated randomisation sequence, with details of the algorithm or programme used), and what was randomised (e.g. treatment to experimental unit, order of treatment for each animal). The EDA has a dedicated feature for randomisation and allocation concealment [18].

#### Examples

1. *“Fifty 12-week-old male Sprague-Dawley rats, weighing 320–360g, were obtained from Guangdong Medical Laboratory Animal Center (Guangzhou, China) and randomly divided into two groups (25 rats/group): the intact group and the castration group. Random numbers were generated using the standard = RAND() function in Microsoft Excel.” [65]*
2. *“Animals were randomized after surviving the initial I/R, using a computer based random order generator.” [66]*
3. *“At each institute, phenotyping data from both sexes is collected at regular intervals on age-matched wildtype mice of equivalent genetic backgrounds. Cohorts of at least seven homozygote mice of each sex per pipeline were generated…. The random allocation of mice to experimental group (wildtype versus knockout) was driven by Mendelian Inheritance.” [28]*

4b. To minimise potential confounding factors such as the order of treatments and measurements, or animal/cage location.

#### Explanation

Ensuring there is no systematic difference between animals in different groups apart from the experimental exposure is an important principle throughout the conduct of the experiment. Identifying nuisance variables (sources of variability or conditions that could potentially bias results), and managing them in the design of the experiment increases the sensitivity of the experiment. For example, rodents in cages at the top of the rack may be exposed to higher light levels, which can affect stress [67]. Mitigation strategies for nuisance variables include randomising or counterbalancing the position of animal cages on the rack, and taking measurements or processing samples in a random order (preferably with the investigator blinded to the treatment received; see **item 5 – Blinding**). Such practices help avoid introducing unintentional systematic differences between comparison groups, also known as order effects. Strategies to avoid order effects include counterbalancing, randomising order of treatment, and blocking (see Box 4).

##### Box 4: Considerations for randomisation

**<u>Simple randomisation</u>**

All animals/samples are simultaneously randomised to the treatment groups without considering any other variable. This strategy is rarely appropriate as it cannot ensure that comparison groups are balanced for factors or covariates that might influence the result of an experiment.

**<u>Randomisation within blocks</u>**

Blocking is a method of controlling natural variation among experimental units. This splits up the experiment into smaller sub-experiments (blocks), and treatments are randomised to experimental units within each block [24, 68, 69]. This takes into account nuisance variables that could potentially bias the results (e.g. cage location, day or week of procedure).

Stratified randomisation uses the same principle as randomisation within blocks, only the strata tend to be traits of the animal that are likely to be associated with the response (e.g. weight class or tumour size class). This can lead to differences in the practical implementation of stratified randomisation as compared to block randomisation (e.g. there may not be equal numbers of experimental units in each weight class).

**<u>Other randomisation techniques</u>**

Minimisation is an alternative strategy to allocate animals/samples to treatment group to balance confounding variables that might influence the result of an experiment. With minimisation the treatment allocated to the next animal/sample depends on the characteristics of those animals/samples already assigned. The aim is that each allocation should minimise the imbalance across multiple factors [70]. This approach works well for a continuous nuisance variable such as body weight or starting tumour volume.

**<u>Examples of nuisance variables that can be accounted for in the randomisation strategy</u>**

- Time or day of the experiment
- Litter, cage or fish tank
- Investigator or surgeon – different level of experience in the people administering the treatments, performing the surgeries, or assessing the results may result in varying stress levels in the animals or duration of anaesthesia
- Equipment (e.g. PCR machine, spectrophotometer) – calibration may vary
- Measurement of a study parameter (e.g. initial tumour volume)
- Animal characteristics – sex, age class, weight class
- Location – exposure to light, ventilation and disturbances may vary in cages located at different height or on different racks, which may affect important physiological processes

**<u>Implication for the analysis</u>**

If blocking factors are used in the randomisation, they should also be included in the analysis. Nuisance variables increase variability in the sample, which reduces statistical power. Including a nuisance variable as a blocking factor in the analysis accounts for that variability and can increase the power, thus increasing the ability to detect a real effect with fewer experimental units. However, blocking uses up degrees of freedom and thus reduces the power if the nuisance variable does not have a substantial impact on variability.

Report the methods used to minimise confounding factors alongside the methods used to allocate animals to groups.

#### Examples

1. *“Randomisation was carried out as follows. On arrival from El-Nile Company, animals were assigned a group designation and weighed. A total number of 32 animals were divided into four different weight groups (eight animals per group). Each animal was assigned a temporary random number within the weight range group. On the basis of their position on the rack, cages were given a numerical designation. For each group, a cage was selected randomly from the pool of all cages. Two animals were removed from each weight range group and given their permanent numerical designation in the cages. Then, the cages were randomized within the exposure group.” [71]*
2. *“…test time was between 08.30am to 12.30pm and testing order was randomized daily, with each animal tested at a different time each test day.” [72]*
3. *“Bulls were blocked by BW into four blocks of 905 animals with similar BW and then within each block, bulls were randomly assigned to one of four experimental treatments in a completely randomized block design resulting in 905 animals per treatment. Animals were allocated to 20 pens (181 animals per pen and five pens per treatment).” [73]*

### Item 5. Blinding

Describe who was aware of the group allocation at the different stages of the experiment (during the allocation, the conduct of the experiment, the outcome assessment, and the data analysis).

#### Explanation

Researchers often expect a particular outcome, and can unintentionally influence the experiment or interpret the data in such a way as to support their preferred hypothesis [74]. Blinding is a strategy used to minimise these subjective biases.

Whilst there is primary evidence of the impact of blinding in the clinical literature that directly compares blinded vs unblinded assessment of outcomes [75], there is limited empirical evidence in animal research [76, 77]. There are, however, compelling data from systematic reviews showing that non-blinded outcome assessment leads to the treatment effects being overestimated, and the lack of bias-reducing measures such as randomisation and blinding can contribute to as much as 30-45% inflation of effect sizes [60, 78, 79].

Ideally, investigators should be unaware of the treatment(s) animals have received or will be receiving, from the start of the experiment until the data have been analysed. If this is not possible for all the stages of an experiment (see Box 5), it should always be possible to conduct at least some of the stages blind. This has implications for the organisation of the experiment and may require help from additional personnel, for example a surgeon to perform interventions, a technician to code the treatment syringes for each animal, or a colleague to code the treatment groups for the analysis. Online resources are available to facilitate allocation concealment and blinding [18].

##### Box 5: Blinding during different stages of an experiment

**<u>During allocation</u>**

Allocation concealment refers to concealing the treatment to be allocated to each individual animal from those assigning the animals to groups, until the time of assignment. Together with randomisation, allocation concealment helps minimise selection bias, which can introduce systematic differences between treatment groups.

**<u>During the conduct of the experiment</u>**

Where possible, animal care staff and those who administer treatments should be unaware of allocation groups to ensure that all animals in the experiment are handled, monitored and treated in the same way. Treating different groups differently based on the treatment they have received could alter animal behaviour and physiology, and produce confounds.

Welfare or safety reasons may prevent blinding of animal care staff but in most cases, blinding is possible. For example, if hazardous microorganisms are used, control animals can be considered as dangerous as infected animals. If a welfare issue would only be tolerated for a short time in treated but not control animals, a harm-benefit analysis is needed to decide whether blinding should be used.

**<u>During the outcome assessment</u>**

The person collecting experimental measurements or conducting assessments should not know which treatment each sample/animal received, and which samples/animals are grouped together. Blinding is especially important during outcome assessment, particularly if there is a subjective element (e.g. when assessing behavioural changes or reading histological slides) [76]. Randomising the order of examination can also reduce bias.

If the person assessing the outcome cannot be blinded to the group allocation (e.g. obvious phenotypic or behavioural differences between groups) some, but not all, of the sources of bias could be mitigated by sending data for analysis to a third party, who has no vested interest in the experiment and does not know whether a treatment is expected to improve or worsen the outcome.

**<u>During the data analysis</u>**

The person analysing the data should know which data are grouped together to enable group comparisons, but should not be aware of which treatment each group received. This type of blinding is often neglected, but is important as the analyst makes many semi-subjective decisions such as applying data transformation to outcome measures, choosing methods for handling missing data and handling outliers. How these decisions will be made should also be decided *a priori*.

Data can be coded prior to analysis so that the treatment group cannot be identified before analysis is completed.

Specify whether blinding was used or not for each step of the experimental process (see Box 5). If blinding was not possible during a specific stage of the experiment, provide the reason why.

#### Examples

1. *“For each animal, four different investigators were involved as follows: a first investigator (RB) administered the treatment based on the randomization table. This investigator was the only person aware of the treatment group allocation. A second investigator (SC) was responsible for the anaesthetic procedure, whereas a third investigator (MS, PG, IT) performed the surgical procedure. Finally, a fourth investigator (MAD) (also unaware of treatment) assessed GCPS and NRS, MNT, and sedation NRS scores.” [80]*
2. *“…due to overt behavioral seizure activity the experimenter could not be blinded to whether the animal was injected with pilocarpine or with saline.” [81]*
3. *“Investigators could not be blinded to the mouse strain due to the difference in coat colors, but the three-chamber sociability test was performed with ANY-maze video tracking software (Stoelting, Wood Dale, IL, USA) using an overhead video camera system to automate behavioral testing and provide unbiased data analyses. The one-chamber social interaction test requires manual scoring and was analyzed by an individual with no knowledge of the questions.” [82]*

### Item 6. Outcome measures

6a. Clearly define all outcome measures assessed (e.g. cell death, molecular markers, or behavioural changes).

#### Explanation

An outcome measure (also known as a dependent variable or a response variable) is any variable recorded during a study (e.g. volume of damaged tissue, number of dead cells, specific molecular marker) to assess the effects of a treatment or experimental intervention. Outcome measures may be important for characterising a sample (e.g. baseline data) or for describing complex responses (e.g. ‘haemodynamic’ outcome measures including heart rate, blood pressure, central venous pressure, and cardiac output).

Explicitly describe what was measured, especially when measures can be operationalised in different ways. For example, activity could be recorded as time spent moving or distance travelled. Where possible, the recording of outcome measures should be made in an unbiased manner (e.g. blinded to the treatment allocation of each experimental group; see **item 5 – Blinding**). Specify how the outcome measure(s) assessed are relevant to the objectives of the study.

#### Example

1. *“The following parameters were assessed: threshold pressure (TP; intravesical pressure immediately before micturition); post-void pressure (PVP; intravesical pressure immediately after micturition); peak pressure (PP; highest intravesical pressure during micturition); capacity (CP; volume of saline needed to induce the first micturition); compliance (CO; CP to TP ratio); frequency of voiding contractions (VC) and frequency of non-voiding contractions (NVCs).” [83]*

6b. For hypothesis-testing studies, specify the primary outcome measure, i.e. the outcome measure that was used to determine the sample size.

#### Explanation

In a hypothesis-testing experiment, the primary outcome measure answers the main biological question. It is the outcome of greatest importance, identified in the planning stages of the experiment and used as the basis for the sample size calculation (see Box 3). For exploratory studies it is not necessary to identify a single primary outcome and often multiple outcomes are assessed (see **item 13 – Objectives**).

In a hypothesis-testing study powered to detect an effect on the primary outcome measure, data on secondary outcomes are used to evaluate additional effects of the intervention but subsequent statistical analysis of secondary outcome measures may be underpowered, making results and interpretation less reliable [84, 85]. Studies that claim to test a hypothesis but do not specify a pre-defined primary outcome measure, or those that change the primary outcome measure after data were collected (also known as primary outcome switching) are liable to selectively report only statistically significant results, favouring more positive findings [86].

Registering a protocol in advance protects the researcher against concerns about selective outcome reporting (also known as data dredging or p-hacking) and provides evidence that the primary outcome reported in the manuscript accurately reflects what was planned [87] (see **item 19 – Protocol registration**).

If the study was designed to test a hypothesis and more than one outcome was assessed, explicitly identify the primary outcome measure and state if it was defined as such prior to data collection.

#### Examples

1. *“The primary outcome of this study will be forelimb function assessed with the staircase test. Secondary outcomes constitute Rotarod performance, stroke volume (quantified on MR imaging or brain sections, respectively), diffusion tensor imaging (DTI) connectome mapping, and histological analyses to measure neuronal and microglial densities, and phagocytic activity… The study is designed with 80% power to detect a relative 25% difference in pellet-reaching performance in the Staircase test.” [88]*
2. *“The primary endpoint of this study was defined as left ventricular ejection fraction (EF) at the end of follow-up, measured by magnetic resonance imaging (MRI). Secondary endpoints were left ventricular end diastolic volume and left ventricular end systolic volume (EDV and ESV) measured by MRI, infarct size measured by ex vivo gross macroscopy after incubation with triphenyltetrazolium chloride (TTC) and late gadolinium enhancement (LGE) MRI, functional parameters serially measured by pressure volume (PV-)loop and echocardiography, coronary microvascular function by intracoronary pressure- and flow measurements and vascular density and fibrosis on histology. Based on a power calculation (estimated effect 7.5% [6], standard deviation of 5%, a power of 0.9 and alpha of 0.05) 8 pigs per group were needed.” [66]*

### Item 7. Statistical methods

7a. Provide details of the statistical methods used for each analysis.

#### Explanation

In hypothesis-testing studies comparing two or more groups, inferential statistics are used to estimate the size of the effect and to determine the weight of evidence against the null hypothesis. The effect size is the magnitude of the difference between two groups. The description of the statistical analysis should provide enough detail so that another researcher could re-analyse the raw data using the same method and obtain the same results. Relevant information includes what the outcome measures and independent variables were, what statistical analyses were performed, what tests were used to check assumptions, and any data transformations [89]. Give details of any confounders, blocking factors or covariates taken into account for each statistical test, include how the effects of each were mitigated. This allows readers to assess if analysis methods were appropriate.

In exploratory studies where no specific hypothesis was tested, descriptive statistics can be used to summarise the data (see **item 10 – Results**). They do not allow conclusions beyond the data but are important for generating new hypotheses that may be tested in subsequent experiments.

For any study reporting descriptive statistics, explicitly state which measure of central tendency is reported (e.g. mean or median) and which measure of variability is reported (e.g. standard deviation, range, quartiles or interquartile range).

#### Examples

1. *“Analysis of variance was performed using the GLM procedure of SAS (SAS Inst., Cary, NC). Average pen values were used as the experimental unit for the performance parameters. The model considered the effects of block and dietary treatment (5 diets). Data were adjusted by the covariant of initial body weight. Orthogonal contrasts were used to test the effects of SDPP processing (UV vs no UV) and dietary SDPP level (3% vs 6%). Results are presented as least squares means. The level of significance was set at P < 0.05.” [90]*
2. *“All risk factors of interest were investigated in a single model. Logistic regression allows blocking factors and explicitly investigates the effect of each independent variable controlling for the effects of all others…. As we were interested in husbandry and environmental effects, we blocked the analysis by important biological variables (age; backstrain; inbreeding; sex; breeding status) to control for their effect. (The role of these biological variables in barbering behavior, particularly with reference to barbering as a model for the human disorder trichotillomania, is described elsewhere: Garner et al., 2004). We also blocked by room to control for the effect of unknown environmental variables associated with this design variable. We tested for the effect of the following husbandry and environmental risk factors: cage mate relationships (i.e. siblings, non-siblings, or mixed); cage type (i.e. plastic or steel); cage height from floor; cage horizontal position (whether the cage was on the side or the middle of a rack); stocking density; and the number of adults in the cage. Cage material by cage height from floor; and cage material by cage horizontal position interactions were examined, and then removed from the model as they were nonsignificant. N = 1959 mice were included in this analysis” [91]*

7b. Specify the experimental unit that was used for each statistical test.

#### Explanation

Incorrect identification of the experimental unit can lead to pseudoreplication and underpowered studies (see **item 1 – Study design**). For example, measurements from 50 individual cells from a single mouse represent N = 1 when the experimental unit is the mouse. The 50 measurements are subsamples and provide an estimate of measurement error so should be averaged or used in a nested analysis. Reporting N = 50 in this case is an example of pseudoreplication [22]. It underestimates the true variability in a study, which can lead to false positives. If, however, each cell taken from the mouse is then randomly allocated to different treatments and assessed individually, the cell might be regarded as the experimental unit.

Explicitly report the experimental unit used in each statistical analysis.

#### Examples

1. *“For each test, the experimental unit was an individual animal.”[92]*
2. *“Maternal data regarding body weight, food intake, water consumption, urinary cotinine level and hormonal analysis were analyzed using the individual animal as the experimental unit. The data for offspring regarding body weight, food intake, organ weight at necropsy, urinary cotinine level, immunohistochemical cellular distribution, TUNEL+ cells, RT-PCR and hormonal analysis were analyzed using the litter as the experimental unit.” [93]*

7c. Describe any methods used to assess whether the data met the assumptions of the statistical approach.

#### Explanation

Hypothesis tests are based on assumptions about the underlying data. Describing how assumptions were assessed, and whether these assumptions are met by the data, enables readers to assess the suitability of the statistical approach used. If the assumptions are incorrect, the conclusions may not be valid. For example, the assumptions for data used in parametric tests (such as a t-test, Z-test, ANOVA, Pearson’s r coefficient, etc.) are that the data are continuous, the residuals from the analysis are normally distributed, the responses are independent, and that different groups should have similar variances.

There are various tests for normality, for example the Shapiro-Wilk and Kolmogorov-Smirnov tests. However, these tests have to be used cautiously. If the sample size is small, they will struggle to detect non-normality, if the sample size is large, the tests will detect minor deviations. An alternative approach is to evaluate data with visual plots e.g. normal probability plots, box plots, scatterplots. If the residuals of the analysis are not normally distributed, the assumption may be satisfied using a data transformation where the same mathematical function is applied to all data points to produce normally distributed data (e.g. log_e_, log_10_, square root, arcsine).

Other types of outcome measures (binary, categorical, or ordinal) will require different methods of analysis, and each will have different sets of assumptions. For example, categorical data are summarised by counts and percentages or proportions, and are analysed by tests of proportions; these analysis methods assume that data are binary, ordinal or nominal, and independent [94].

Report the type of outcome measure and the methods used to test the assumptions of the statistical approach. If data were transformed, identify precisely the transformation used and which outcome measures it was applied to.

#### Examples

1. *“Model assumptions were checked using the Shapiro-Wilk normality test and Levene’s Test for homogeneity of variance and by visual inspection of residual and fitted value plots. Some of the response variables had to be transformed by applying the natural logarithm or the second or third root, but were back-transformed for visualization of significant effects.” [95]*
2. *The effects of housing (treatment) and day of euthanasia on cortisol levels were assessed by using fixed-effects 2-way ANOVA. An initial exploratory analysis indicated that groups with higher average cortisol levels also had greater variation in this response variable. To make the variation more uniform, we used a logarithmic transform of each fish’s cortisol per unit weight as the dependent variable in our analyses. This action made the assumptions of normality and homoscedasticity (standard deviations were equal) of our analyses reasonable. [96]*

### Item 8. Experimental animals

8a. Provide details of the animals used, including species, strain and substrain, sex, age or developmental stage, and weight.

#### Explanation

The species, strain, substrain, sex, weight, and age of animals are critical factors that can influence most experimental results [97–101]. Reporting the characteristics of all animals used is equivalent to standardised human patient demographic data; these data support both the internal and external validity of the study results. It enables other researchers to repeat the experiment and generalise the findings. It also enables readers to assess whether the animal characteristics chosen for the experiment are relevant to the research objectives.

Report age and weight for each group, include summary statistics (e.g. mean and standard deviation) and, if possible, baseline values for individual animals (e.g. as supplementary information or a link to a publicly accessible data repository). For most species, precise reporting of age is more informative than a description of the developmental status (e.g. a mouse referred to as an adult can vary in age from six to 20 weeks [102]). In some cases, however, reporting the developmental stage is more informative than chronological age, for example in juvenile *Xenopus*, where rate of development can be manipulated by incubation temperature [103].

#### Example

1. *“One hundred and nineteen male mice were used: C57BL/6OlaHsd mice (n = 59), and BALB/c OlaHsd mice (n = 60) (both from Harlan, Horst, The Netherlands). At the time of the EPM test the mice were 13 weeks old and had body weights of 27.4 ± 0.4 g and 27.8 ± 0.3 g, respectively (mean ± SEM).” [104]*
2. *“Histone Methylation Profiles and the Transcriptome of X. tropicalis Gastrula Embryos. To generate epigenetic profiles, ChIP was performed using specific antibodies against trimethylated H3K4 and H3K27 in Xenopus gastrula-stage embryos (Nieuwkoop-Faber stage 11–12), followed by deep sequencing (ChIP-seq). In addition, polyA-selected RNA (stages 10–13) was reverse transcribed and sequenced (RNA-seq).” [105]*

8b. Provide further relevant information on the provenance of animals, health/immune status, genetic modification status, genotype, and any previous procedures.

#### Explanation

The animals’ provenance, their health or immune status and their history of previous testing or procedures, can influence their physiology and behaviour as well as their response to treatments, and thus impact on study outcomes. For example, animals of the same strain, but from different sources, or animals obtained from the same source but at different times, may be genetically different [17]. The immune or microbiological status of the animals can also influence welfare, experimental variability and scientific outcomes [106–108].

Report the health status of animals in the study, and any previous procedures the animals have undergone. For genetically modified animals, describe the genetic modification status (e.g. knockout, overexpression), genotype (e.g. homozygous, heterozygous), manipulated gene(s), genetic methods and technologies used to generate the animals, how the genetic modification was confirmed, and details of animals used as controls (e.g. littermate controls [109]).

Reporting the correct nomenclature is crucial to understanding the data and ensuring that the research is discoverable and replicable [110–112]. Useful resources for reporting nomenclature for different species include:

- Mice - International Committee on Standardized Genetic Nomenclature (https://www.jax.org/jax-mice-and-services/customer-support/technical-support/genetics-and-nomenclature)
- Rats - Rat Genome and Nomenclature Committee (https://rgd.mcw.edu/)
- Zebrafish - Zebrafish Information Network (http://zfin.org/)
- Xenopus - Xenbase (http://www.xenbase.org/entry/)
- Drosophila – FlyBase (http://flybase.org/)

#### Examples

1. *“A construct was engineered for knockin of the miR-128 (miR-128-3p) gene into the Rosa26 locus. Rosa26 genomic DNA fragments (~1.1 kb and ~4.3kb 5’ and 3’ homology arms, respectively) were amplified from C57BL/6 BAC DNA, cloned into the pBasicLNeoL vector sequentially by in-fusion cloning, and confirmed by sequencing. The miR-128 gene, under the control of tetO-minimum promoter, was also cloned into the vector between the two homology arms. In addition, the targeting construct also contained a loxP sites flanking the neomycin resistance gene cassette for positive selection and a diphtheria toxin A (DTA) cassette for negative selection. The construct was linearized with ClaI and electroporated into C57BL/6N ES cells. After G418 selection, seven-positive clones were identified from 121 G418-resistant clones by PCR screening. Six-positive clones were expanded and further analyzed by Southern blot analysis, among which four clones were confirmed with correct targeting with single-copy integration. Correctly targeted ES cell clones were injected into blastocysts, and the blastocysts were implanted into pseudo-pregnant mice to generate chimeras by Cyagen Biosciences Inc. Chimeric males were bred with Cre deleted mice from Jackson Laboratories to generate neomycin-free knockin mice. The correct insertion of the miR-128 cassette and successful removal of the neomycin cassette were confirmed by PCR analysis with the primers listed in Supplementary Table 1.” [113]*
2. *“The C57BL/6J (Jackson) mice were supplied by Charles River Laboratories. The C57BL/6JOlaHsd (Harlan) mice were supplied by Harlan. The α-synuclein knockout mice were kindly supplied by Prof. [X] (Cardiff University, Cardiff, United Kingdom.) and were congenic C57BL/6JCrl (backcrossed for 12 generations). TNFα−/− mice were kindly supplied by Dr. [Y] (Queens University, Belfast, Northern Ireland) and were inbred on a homozygous C57BL/6J strain originally sourced from Bantin & Kingman and generated by targeting C57BL/6 ES cells. T286A mice were obtained from Prof. [Z] (University of California, Los Angeles, CA). These mice were originally congenic C57BL/6J (backcrossed for five generations) and were then inbred (cousin matings) over 14 y, during which time they were outbred with C57BL/6JOlaHsd mice on three separate occasions.” [114]*

### Item 9. Experimental procedures

For each experimental group, including controls, describe the procedures in enough detail to allow others to replicate them, including:

9a. What was done, how it was done and what was used.

#### Explanation

Essential information to describe in the manuscript includes the procedures used to develop the model (e.g. induction of the pathology), the procedures used to measure the outcomes, and pre- and post-experimental procedures, including animal handling and welfare monitoring. Animal handling can be a source of stress and the specific method used (e.g. mice picked up by tail or in cupped hands) can affect research outcomes [56, 115, 116]. Details about animal care and monitoring intrinsic to the procedure are discussed in further detail in **item 16 – Animal care and monitoring**. Provide enough detail to enable others to replicate the methods and highlight any quality assurance and quality control used [117, 118]. A schematic of the experimental procedures with a timeline can give a clear overview of how the study was conducted. Information relevant to distinct types of interventions and resources are described in Box 6.

##### Box 6: Examples of information to include when reporting specific types experimental procedures and resources

| Procedures | Resources |
| --- | --- |
| <u>Pharmacological procedures (intervention and control)</u><br>Drug formulation<br>Dose<br>Volume<br>Concentration<br>Site and route of administration<br>Frequency of administration<br>Vehicle or carrier solution formulation and volume<br>Any evidence that the pharmacological agent used reaches the target tissue<br><u>Surgical procedures (including sham surgery)</u><br>Description of the surgical procedure<br>Anaesthetic used (including dose and other information listed in pharmacological procedures section above)<br>Pre and post analgesia regimen<br>Pre-surgery procedures (e.g. fasting)<br>Aseptic techniques<br>Monitoring (e.g. assessment of surgical anaesthetic plane) | <u>Cell lines</u><br>Identification<br>Provenance<br>Verification and authentication<br>RRID [120, 121]<br><u>Reagents (e.g. antibodies, chemicals)</u><br>Manufacturer<br>Supplier<br>Catalogue number<br>Lot number (if applicable)<br>RRID<br><u>Equipment and software</u><br>Manufacturer<br>Supplier<br>Model/version number<br>Calibration procedures (if applicable)<br>RRID |
| Whether the procedure is terminal or not<br>Post-surgery procedures<br>Duration of the procedure and duration of anaesthesia<br>Physical variables measured<br><u>Pathogen infection (intervention and control)</u><br>Infectious agent<br>Dose load<br>Vehicle or carrier solution formulation and volume<br>Site and route of infection<br>Timing or frequency of infection<br><u>Euthanasia</u><br>Method of euthanasia, including the humane standards the method complies with, such as AVMA [119]<br>Pharmacological agent, if used (including dose and information listed in pharmacological procedures section above)<br>Any measures taken to reduce pain and distress before or during euthanasia<br>Timing of euthanasia<br>Tissues collected post-euthanasia and timing of collection |  |

Where available, cite the Research Resource Identifier (RRID) for reagents and tools used [120, 121]. RRIDs are unique and stable, allowing unambiguous identification of reagents or tools used in a study, aiding other researchers to replicate the methods.

Detailed step-by-step procedures can also be saved and shared online, for example using Protocols.io [122], which assigns a DOI to the protocol and allows cross-referencing between protocols and publications.

#### Examples

1. *“Fig… shows the timeline for instrumentation, stabilization, shock/injury, and resuscitation…. Animals were food-deprived for 18 hours before surgery, but allowed free access to water. On the morning of surgery, swine were sedated with tiletamine-zolazepam (Telazol^®^; 5-8 mg/kg IM; Zoetis Inc., Kalamazoo MI) in the holding pen, weighed, and masked with isoflurane (3%, balance 100% O2) for transport to the lab. The marginal ear vein was catheterized for administration of atropine (0.02 mg/kg IV; Sparhawk Laboratories, Lenexa KS), and buprenorphine for pre-emptive analgesia (3 mg/ml IV; ZooPharm, Laramie WY). Ophthalmic ointment (Puralube^®^; Fera Pharmaceuticals) was applied to prevent corneal drying. Animals were intubated in dorsal recumbency with a cuffed 6 or 7 Fr endotracheal tube. Anesthetic plane was maintained by isoflurane (1-1.5%; 21-23 % O2, balance N2). Oxygen saturation (sPO2, %) and heart rate (HR) were monitored with a veterinary pulse oximeter placed in the buccal cavity (Masimo SET Radical-7; Irvine CA). Core temperature was monitored with a rectal probe and maintained at 36.5-38°C with a microprocessor-controlled feedback water blanket (Blanketrol^®^ II, Cincinnati Sub-Zero (CSZ) Cincinnati, OH) placed under the animal. Anesthetic depth was assessed every 5 min for the duration of the experiment by reflexes (corneal touch, pedal flexion, coronary band pinch) and vital signs (sPO2, HR, core temperature).* *Fig…. Experimental time line for instrumentation, shock/injury, resuscitation, post-resuscitation monitoring, and blood sampling in a swine model of polytrauma and coagulopathy.” [123]*
2. *“For the diet-induced obesity (DIO) model, eight-week-old male mice had ad libitum access to drinking water and were kept on standard chow (SFD, 10.9 kJ/g) or on western high-fat diet (HFD; 22 kJ/g; kcal from 42% fat, 43% from carbohydrates and 15% from protein; E15721-34, Ssniff, Soest, Germany) for 15 weeks (http://dx.doi.org/10.17504/protocols.io.kbacsie).” [124]*

9b. When and how often.

#### Explanation

Clearly report the frequency and timing of experimental procedures and measurements, including the light and dark cycle (e.g. 12:12), circadian time cues (e.g. lights on at 08:00), and experimental time sequence (e.g. interval between baseline and comparator measurements or interval between procedures and measurements). Along with innate circadian rhythms, these can affect research outcomes such as behavioural, physiological, and immunological parameters [125, 126]. Also report the timing and frequency of welfare assessments, taking into consideration the normal activity patterns (see **item 16 – Animal care and monitoring**). For example, nocturnal animals may not show behavioural signs of discomfort during the day [127].

#### Examples

1. *“Blood pressure, heart rate, oxygen saturation and amount of blood extracted were recorded every 5 minutes. Blood samples were drawn at baseline (pre injury), 0 minutes (immediately after injury), and after 30 and 60 minutes.” [128]*
2. *“After a 5-h fast (7:30–12:30am), awake and freely moving mice were randomized and subjected to three consecutive clamps performed in the same mice as described above, with a 2 days recovery after each hyperinsulinemic/hypoglycemic (mHypo, n = 6) or hyperinsulinemic/euglycemic (mEugly, n = 4) clamps.” [129]*

9c. Where (including detail of any acclimation periods).

#### Explanation

Physiological acclimation after a stressful event, such as transport (e.g. between supplier, animal facility and laboratory), but before the experiment begins allows stabilisation of physiological responses of the animal [130, 131]. Protocols vary depending on species, strain, and outcome; for example physiological acclimation following transportation of different animals can take anywhere from 24 hours to more than one week [132]. Procedural acclimation, immediately before a procedure, allows stabilisation of the animals’ responses after unaccustomed handling, novel environments, and previous procedures, which otherwise can induce behavioural and physiological changes [133, 134].

Indicate where studies were performed (e.g. dedicated laboratory space or animal facility, home cage, open field arena, water maze) and whether periods of physiological or procedural acclimation were included in the study protocol, including type and duration. If the study involved multiple sites explicitly state where each experiment and sample analysis was performed. Include any accreditation of laboratories if appropriate (e.g. if samples are sent to a commercial laboratory for analysis).

#### Example

1. *“Fish were singly housed for 1 week before being habituated to the conditioning tank over 2 consecutive days. The conditioning tank consisted of an opaque tank measuring 20 cm (w) 15 cm (h) 30 cm (l) containing 2.5 l of aquarium water with distinct visual cues (spots or stripes) on walls at each end of the tank… During habituation, each individual fish was placed in the conditioning apparatus for 20 minutes with free access to both compartments and then returned to its home tank.” [135]*

9d. Why (provide rationale for procedures).

#### Explanation

There may be numerous approaches to investigate any given research problem, therefore it is important to explain why a particular procedure or technique was chosen. This is especially relevant when procedures are novel or specific to a research laboratory, or constrained by the animal model or experimental equipment (e.g. route of administration determined by animal size [136]).

#### Examples

1. *“Because of the very small caliber of the murine tail veins, partial paravenous injection is common if 18F-FDG is administered by tail vein injection (intravenous). This could have significantly biased our comparison of the biodistribution of 18F-FDG under various conditions. Therefore, we used intraperitoneal injection of 18F-FDG for our experiments evaluating the influence of animal handling on 18F-FDG biodistribution.” [137]*
2. *“Since Xenopus oocytes have a higher potential for homologous recombination than fertilized embryos (Hagmann et al., 1996), we next tested whether the host transfer method could be used for efficient HDR-mediated knock-in. We targeted the C-terminus of X. laevis Ctnnb1 (β-catenin), a key cytoskeletal protein and effector of the canonical Wnt pathway, because previous studies have shown that addition of epitope tags to the C-terminus do not affect the function of the resulting fusion protein (Fig. 2A) (Evans et al., 2010; Miller and Moon, 1997). CRISPR components were injected into X. laevis oocytes followed by host transfer or into embryos.” [138]*

### Item 10. Results

For each experiment conducted, including independent replications, report:

10a. Summary/descriptive statistics for each experimental group, with a measure of variability where applicable.

#### Explanation

Summary/descriptive statistics provide a quick and simple description of the data, they communicate quantitative results easily and facilitate visual presentation. For continuous data, these descriptors include a measure of central tendency (e.g. mean, median) and a measure of variability (e.g. quartiles, range, standard deviation) to help readers assess the precision of the data collected. Categorical data can be expressed as counts, frequencies, or proportions.

Report data for all experiments conducted. If a complete experiment is repeated on a different day, or under different conditions, report the results of all repeats, rather than selecting data from representative experiments. Report the exact number of experimental units per group so readers can gauge reliability of results (see **item 2 – Sample size**, and **item 3 – Inclusion and exclusion criteria**). Present data clearly as text, in tables, or in graphs, to enable information to be evaluated, or extracted for future meta-analyses. Report descriptive statistics with a clearly identified measure of variability for each group. Example 1 shows data summarised as medians and, in brackets, lower and upper quartiles. Boxplots are a convenient way to summarise continuous data, plotted as median, interquartile range, minimum, maximum and outliers, as shown in Example 2.

#### Examples

1. *“Features of autism compared between the control group and the autistic model groups” [139]*
2. *“Boxplots of median heart rate (beats per minute, bpm) during rs-fMRI scans for Y (n = 12), AU (n = 12), and AI (n = 12) animals.”[140]*

|  | Median (quartiles) |  |
| --- | --- | --- |
|  | Control group (n=13) | Autism model group (n=14) |
| Birth weight (g) | 4.2 (3.9 – 4.5) | 3.6 (2.9 – 5.0) |
| Weight at 28 weeks (g) | 47.8 (44.0 – 51.8) | 32.5 (29.0 – 53.5) |
| Number of squares traversed | 6.0 (4.0 – 11.0) | 3.0 (2.0 – 7.0) |
| Rearing movements | 6.0 (4.0 – 13.0) | 3.0 (2.0 – 6.0) |
| Grooming movements (number of rats performing grooming movements/total in the group) | 4/13 | 2/14 |

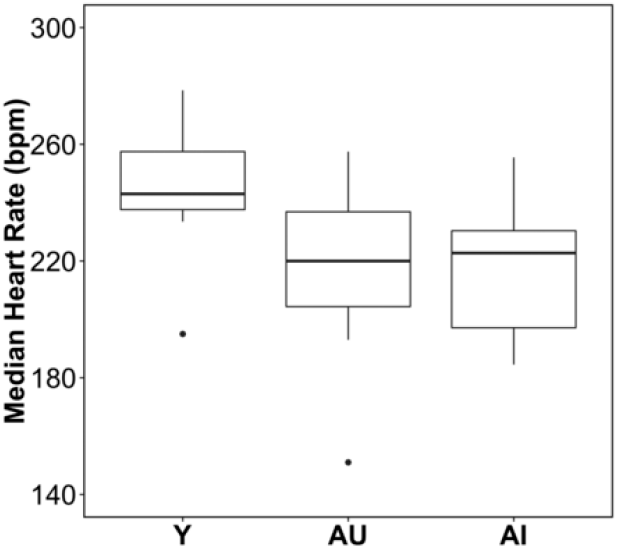

10b. If applicable, the effect size with a confidence interval.

#### Explanation

An effect size is a quantitative measure that estimates the magnitude of differences between groups, or relationships between variables. It can be used to assess the patterns in the data collected and make inferences about the wider population from which the sample came. The confidence interval for the effect indicates how precisely the effect has been estimated, and tells the reader about the strength of the effect [141]. For example, if zero is included in the 95% confidence interval, the presence of an effect cannot be assumed. In studies where statistical power is low, and/or hypothesis-testing is inappropriate, providing the effect size and confidence interval indicates how small or large an effect might really be, so a reader can judge the biological significance of the data [142, 143]. Reporting effect sizes with confidence intervals also facilitates extraction of useful data for systematic review and meta-analysis. Where multiple independent studies included in a meta-analysis show quantitatively similar effects, even if each is statistically non-significant, this provides powerful evidence that a relationship is ‘real’, although small.

Report all analyses performed, even those providing non-statistically significant results. Report the effect size, to indicate the size of the difference between groups in the study, with a confidence interval, to indicate the precision of the effect size estimate.

#### Example

1. *“For all traits identified as having a significant genotype effect for the Usp47tm1b(EUCOMM)Wtsi line (MGI:5605792), a comparison is presented of the standardized genotype effect with 95% confidence interval for each sex with no multiple comparisons correction. Standardization, to allow comparison across variables, was achieved by dividing the genotype estimate by the signal seen in the wildtype population. Shown in red are statistically significant estimates. RBC: red blood cells; BMC: bone mineral content; BMD: bone mineral density; WBC: white blood cells.” [28]*

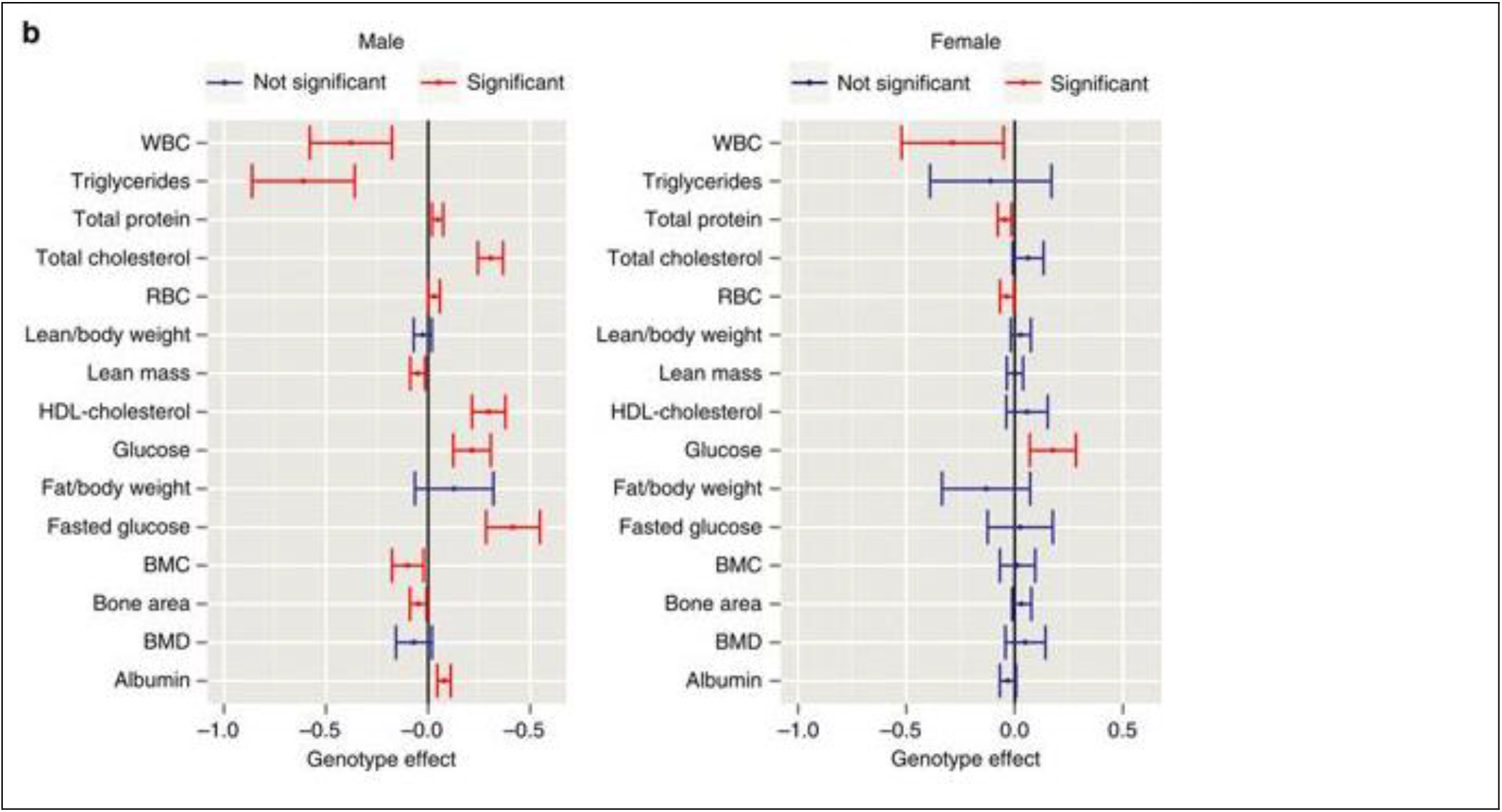

## 2. Recommended Set

The Recommended Set (Box 7) adds context to the study described, including further detail about the methodology and advice on what to include in the more narrative parts of a manuscript. Items are presented in a logical order, there is no ranking within the set.

### Box 7: ARRIVE Recommended Set

11. Abstract
12. Background
13. Objectives
14. Ethical statement
15. Housing and husbandry
16. Animal care and monitoring
17. Interpretation /scientific implications
18. Generalisability /translation
19. Protocol registration
20. Data access
21. Declaration of interests

### Item 11. Abstract

Provide an accurate summary of the research objectives, animal species, strain and sex, key methods, principal findings, and study conclusions.

#### Explanation

A transparent and accurate abstract increases the utility and impact of the manuscript, and allows readers to assess the reliability of the study [144]. The abstract is often used as a screening tool by readers to decide whether to read the full article or to select an article for inclusion in a systematic review. However, abstracts often either do not contain enough information for this purpose [145, 146], or contain information that is inconsistent with the results in the rest of the manuscript [147, 148].

To maximise utility, include details of the species, sex and strain of animals used, and accurately report the methods, results and conclusions of the study. Also describe the objectives of the study, including whether it was designed to either test a specific hypothesis or to generate a new hypothesis (see **Item 13 – Objectives**). Incorporating this information will enable readers to interpret the strength of evidence, and judge how the study fits within the wider knowledge base.

#### Examples

1. *“BACKGROUND AND PURPOSE: Asthma is an inflammatory disease that involves airway hyperresponsiveness and remodelling. Flavonoids have been associated to anti-inflammatory and antioxidant activities and may represent a potential therapeutic treatment of asthma. Our aim was to evaluate the effects of the sakuranetin treatment in several aspects of experimental asthma model in mice.* *EXPERIMENTAL APPROACH: Male BALB/c mice received ovalbumin (i.p.) on days 0 and 14, and were challenged with aerolized ovalbumin 1% on days 24, 26 and 28. Ovalbumin-sensitized animals received vehicle (saline and dimethyl sulfoxide, DMSO), sakuranetin (20 mg kg–1per mice) or dexamethasone (5 mg kg–1 per mice) daily beginning from 24th to29th day. Control group received saline inhalation and nasal drop vehicle. On day 29, we determined the airway hyperresponsiveness, inflammation and remodelling as well as specific IgE antibody. RANTES, IL-5, IL-4, Eotaxin, IL-10, TNF-a, IFN-g and GMC-SF content in lung homogenate was performed by Bioplex assay, and 8-isoprostane and NF-kB activations were visualized in inflammatory cells by immunohistochemistry*. *KEY RESULTS: We have demonstrated that sakuranetin treatment attenuated airway hyperresponsiveness, inflammation and remodelling; and these effects could be attributed to Th2 pro-inflammatory cytokines and oxidative stress reduction as well as control of NF-kB activation*. *CONCLUSIONS AND IMPLICATIONS: These results highlighted the importance of counteracting oxidative stress by flavonoids in this asthma model and suggest sakuranetin as a potential candidate for studies of treatment of asthma.” [149]*
2. *“In some parts of the world, the laboratory pig (Sus scrofa) is often housed in individual, sterile housing which may impose stress. Our objectives were to determine the effects of isolation and enrichment on pigs housed within the PigTurn^®^ — a novel penning system with automated blood sampling — and to investigate tear staining as a novel welfare indicator. Twenty Yorkshire × Landrace weaner pigs were randomly assigned to one of four treatments in a 2 × 2 factorial combination of enrichment (non-enriched [NE] or enriched [E]) and isolation (visually isolated [I] or able to see another pig [NI]). Pigs were catheterised and placed into the PigTurns^®^ 48 h post recovery. Blood was collected automatically twice daily to determine white blood cell (WBC) differential counts and assayed for cortisol. Photographs of the eyes were taken daily and tear staining was quantified using a 0–5 scoring scale and Image-J software to measure stain area and perimeter. Behaviour was video recorded and scan sampled to determine time budgets. Data were analysed as an REML using the MIXED procedure of SAS. Enrichment tended to increase proportion of time standing and lying laterally and decrease plasma cortisol, tear-stain area and perimeter. There was a significant isolation by enrichment interaction. Enrichment given to pigs housed in isolation had no effect on plasma cortisol, but greatly reduced it in non-isolated pigs. Tear-staining area and perimeter were highest in the NE-I treatment compared to the other three treatments. Eosinophil count was highest in the E-NI treatment and lowest in the NE-I treatment. The results suggest that in the absence of enrichment, being able to see another animal but not interact may be frustrating. The combination of no enrichment and isolation maximally impacted tear staining and eosinophil numbers. However, appropriate enrichment coupled with proximity of another pig would appear to improve welfare.” [150]*

### Item 12. Background

12a. Include sufficient scientific background to understand the rationale and context for the study, and explain the experimental approach.

#### Explanation

Scientific background information for an animal study should demonstrate a clear evidence gap and explain why an *in vivo* approach was warranted. Systematic reviews of the animal literature provide the most convincing evidence that a research question has not been conclusively addressed, by showing the extent of current evidence within a field of research. They can also inform the choice of animal model by providing a comprehensive overview of the models used along with their benefits and limitations [151, 152].

Describe the rationale and context of the study and how it relates to other research, including relevant references to previous work. Outline evidence underpinning the hypothesis or objectives and explain why the experimental approach is best suited to answer the research question.

#### Example

1. *“For decades, cardiovascular disease has remained the leading cause of mortality worldwide…[and] cardiovascular research has been performed using healthy and young, non-diseased animal models. Recent failures of cardioprotective therapies in obese insulin-resistant, diabetic, metabolic syndrome-affected and aged animals that were otherwise successful in healthy animal models has highlighted the need for the development of animal models of disease that are representative of human clinical conditions… In the clinical setting, elderly male patients often present with both testosterone deficiency (TD) and the metabolic syndrome (MetS). A strong and compounding association exists between MetS and TD which may have significant impact on cardiovascular disease and its outcomes which is not addressed by current models…. their mutual presentation in the clinical setting warrants the development of appropriate animal models of the MetS with hypogonadism, especially in the context of cardiovascular disease research.” [153]*

12b. Explain how the animal species and model used address the scientific objectives and, where appropriate, the relevance to human biology.

#### Explanation

Provide enough detail for the reader to assess the suitability of the animal model used to address the research question. Include information on the rationale for choosing a particular species, explain how the outcome measures assessed are relevant to the condition under study, and how the model was validated. Stating that an animal model is commonly used in the field is not appropriate, and a well-considered, detailed rationale should be provided.

When the study models a human disease, indicate how the model is appropriate for addressing the specific objectives of the study [154]. This can include a description of how the induction of the disease, disorder, or injury is sufficiently analogous to the human condition, how the model responds to known clinically-effective treatments, how similar symptoms are to the clinical disease and how animal characteristics were selected to represent the age, sex, and health status of the clinical population [14].

#### Examples

1. *“… we selected a pilocarpine model of epilepsy that is characterized by robust, frequent spontaneous seizures acquired after a brain insult, well-described behavioral abnormalities, and poor responses to antiepileptic drugs. These animals recapitulate several key features of human temporal lobe epilepsy, the most common type of epilepsy in adults.” [155]*
2. *“Transplantation of healthy haematopoietic stem cells (HSCs) is a critical therapy for a wide range of malignant haematological and non-malignant disorders and immune dysfunction (Snowden et al., 2012; Sykes & Nikolic, 2005; Thomas et al., 1957)…. Zebrafish are already established as a successful model to study the haematopoietic system, with significant homology with mammals (de Jong & Zon, 2005; Gering & Patient, 2005; Kissa & Herbomel, 2010; Renshaw & Trede, 2012; Traver et al., 2003; White et al., 2008). Imaging of zebrafish transparent embryos remains a powerful tool and has been critical to confirm that the zebrafish Caudal Haematopoietic Tissue (CHT) is comparable to the mammalian foetal haematopoietic niche (Gering & Patient, 2005; Kissa & Herbomel, 2010; Tamplin et al., 2015). Xenotransplantation in zebrafish embryos has revealed highly conserved mechanisms between zebrafish and mammals. Recently, murine bone marrow cells were successfully transplanted into zebrafish embryos, revealing highly conserved mechanism of haematopoiesis between zebrafish and mammals (Parada-Kusz et al., 2017). Additionally, CD34 enriched human cells transplanted into zebrafish were shown to home to the CHT and respond to zebrafish stromal-cell derived factors (Staal et al., 2016).” [156]*

### Item 13. Objectives

Clearly describe the research question, research objectives and, where appropriate, specific hypotheses being tested.

#### Explanation

Explaining the purpose of the study by describing the question(s) that the research addresses, allows readers to determine if the study is relevant to them. Readers can also assess the relevance of the model organism, procedures, outcomes measured, and analysis used.

Knowing if a study is exploratory or hypothesis-testing is critical to its interpretation. A typical exploratory study may measure multiple outcomes and look for patterns in the data, or relationships that can be used to generate hypotheses. It may also be a pilot study which aims to inform the design or feasibility of larger subsequent experiments. Exploratory research helps researchers to design hypothesis-testing experiments, by choosing what variables or outcome measures to focus on in subsequent studies.

Testing a specific hypothesis has implications for both the study design and the data analysis [17, 157]. For example, an experiment designed to detect a hypothesised effect will likely need to be analysed with inferential statistics, and a statistical estimation of the sample size will need to be performed *a priori* (see **item 2 – Sample size**). Hypothesis-testing studies also have a pre-defined primary outcome measure, which is used to assess the evidence in support of the specific research question (see **item 6 – Outcome measures**).

In contrast, exploratory research investigates many possible effects, and is likely to yield more false positive results because some will be positive by chance; thus results from well-designed hypothesis-testing studies provide stronger evidence. Independent replication and meta-analysis can further increase the confidence in conclusions.

Clearly outline the objective(s) of the study, including whether it is hypothesis-testing or exploratory, or if it includes research of both types. Hypothesis-testing studies may collect additional information for exploratory purposes, it is important to distinguish which hypotheses were prespecified and which originated after data inspection, especially when reporting unanticipated effects or outcomes that were not part of the original study design.

#### Examples

1. *“The primary objective of this study was to investigate the cellular immune response to MSC injected into the striatum of allogeneic recipients (6-hydroxydopamine [6-OHDA]-hemilesioned rats, an animal model of Parkinson’s disease [PD]), and the secondary objective was to determine the ability of these cells to prevent nigrostriatal dopamine depletion and associated motor deficits in these animals.” [158]*
2. *“In this exploratory study, we aimed to investigate whether calcium electroporation could initiate an anticancer immune response similar to electrochemotherapy. To this end, we treated immunocompetent balb/c mice with CT26 colon tumors with calcium electroporation, electrochemotherapy, or ultrasound-based delivery of calcium or bleomycin.” [159]*

### Item 14. Ethical statement

Provide the name of the ethical review committee or equivalent that has approved the use of animals in this study and any relevant licence or protocol numbers (if applicable). If ethical approval was not sought or granted, provide a justification.

#### Explanation

Authors are responsible for complying with regulations and guidelines relating to the use of animals for scientific purposes. This includes ensuring that they have the relevant approval for their study from an appropriate ethics committee and/or regulatory body before the work starts. The ethical statement provides editors, reviewers and readers with assurance that studies have received this ethical oversight [12]. This also promotes transparency and understanding about the use of animals in research and fosters public trust.

Provide a clear statement explaining how the study conforms to appropriate regulations and guidelines, and indicate relevant licence/protocol numbers so that the study can be identified. Add also any relevant accreditation e.g. AAALAC (American Association for Accreditation of Laboratory Animal Care) [160] or GLP (Good Laboratory Practice).

If the research is not covered by any regulation and formal ethical approval is not required (e.g. a study using animal species not protected by regulations or law), demonstrate that international standards were complied with and cite the appropriate reference. In such cases, provide a clear statement explaining why the research is exempt from regulatory approval.

#### Examples

1. *“All procedures were conducted in accordance with the United Kingdom Animal (Scientific Procedures) Act 1986, approved by institutional ethical review committees (Alderley Park Animal Welfare and Ethical Review Board and Babraham Institute Animal Welfare and Ethical Review Board) and conducted under the authority of the Project Licence (40/3729 and 70/8307, respectively).” [161]*
2. *“All protocols in this study were approved by the Committee on the Ethics of Animal Experiments of Fuwai Hospital, Peking Union Medical College and the Beijing Council on Animal Care, Beijing, China (IACUC permit number: FW2010-101523), in compliance with the Guide for the Care and Use of Laboratory Animals published by the US National Institutes of Health (NIH publication no.85-23, revised 1996).” [162]*
3. *“Samples and data were collected according to Institut de Sélection d’Animale (ISA) protocols, under the supervision of ISA employees. Samples and data were collected as part of routine animal data collection in a commercial breeding program for layer chickens in The Netherlands. Samples and data were collected on a breeding nucleus of ISA for breeding purposes only, and is a non-experimental, agricultural practice, regulated by the Act Animals, and the Royal Decree on Procedures. The Dutch Experiments on Animals Act does not apply to non-experimental, agricultural practices. An ethical review by the Statement Animal Experiment Committee was therefore not required. No extra animal discomfort was caused for sample collection for the purpose of this study.” [163]*

### Item 15. Housing and husbandry

Provide details of housing and husbandry conditions, including any environmental enrichment.

#### Explanation

The environment determines the health and wellbeing of the animals and every aspect of it can potentially affect their behavioural and physiological responses, thereby affecting research outcomes. Different studies may be sensitive to different environmental factors, and particular aspects of the environment necessary to report may depend on the type of study. Examples of housing and husbandry conditions known to affect animal welfare and research outcomes are listed in Table 1; consider reporting these elements and any other housing and husbandry conditions likely to influence the study outcomes.

**Table 1:**
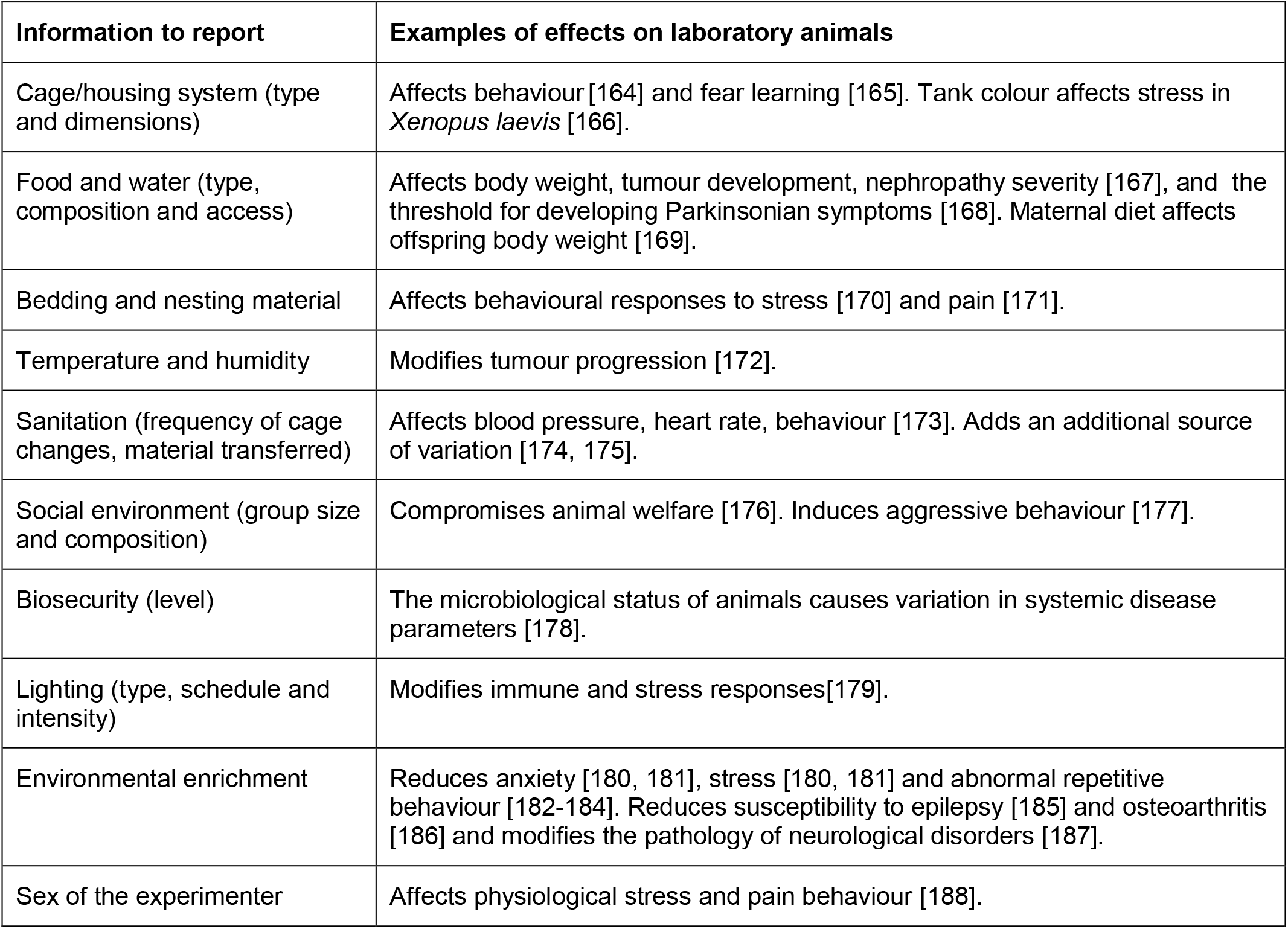
Examples of information to consider when reporting housing and husbandry, and their effects on laboratory animals

Environment, either deprived or enriched, can affect a wide range of physiological and behavioural responses [189]. Specific details to report include, but are not limited to, structural enrichment (e.g. elevated surfaces, dividers), resources for species-typical activities (e.g. nesting material, shelters, gnawing sticks), toys and or other tools used to stimulate exploration, exercise (e.g. running wheel), and novelty. If no environmental enrichment was provided, this should be clearly stated with justification. Similarly, scientific justification needs to be reported for withholding food and water [190], and for singly housing animals [191, 192].

#### Examples

1. *“Breeding colonies were kept in individually ventilated cages (IVCs; Tecniplast, Italy) at a temperature of 20°C to 24°C, humidity of 50% to 60%, 60 air exchanges per hour in the cages, and a 12/12-hour light/dark cycle with the lights on at 5:30 AM. The maximum caging density was five mice from the same litter and sex starting from weaning. As bedding, spruce wood shavings (Lignocel FS-14; J. Rettenmaier und Soehne GmbH, Rosenberg, Germany) were provided. Mice were fed a standardized mouse diet (1314, Altromin, Germany) and provided drinking water ad libitum. All materials, including IVCs, lids, feeders, bottles, bedding, and water were autoclaved before use. Sentinel mice were negative for at least all Federation of laboratory animal science associations (FELASA)-relevant murine infectious agents [30] as diagnosed by our health monitoring laboratory, mfd Diagnostics GmbH, Wendelsheim, Germany.” [193]*
2. *“Same sex litter mates were housed together in individually ventilated cages with two or four mice per cage. All mice were maintained on a regular diurnal lighting cycle (12:12 light:dark) with ad libitum access to food (7012 Harlan Teklad LM-485 Mouse/Rat Sterilizable Diet) and water. Chopped corn cob was used as bedding. Environmental enrichment included nesting material (Nestlets, Ancare, Bellmore, NY, USA), PVC pipe, and shelter (Refuge XKA-2450-087, Ketchum Manufacturing Inc., Brockville, Ontario, Canada). Mice were housed under broken barrier-specific pathogen-free conditions in the Transgenic Mouse Core Facility of Cornell University, accredited by AAALAC (The Association for Assessment and Accreditation of Laboratory Animal Care International).” [194]*

### Item 16. Animal care and monitoring

16a. Describe any interventions or steps taken in the experimental protocols to reduce pain, suffering and distress.

#### Explanation

A safe and effective analgesic plan is critical to relieve pain, suffering and distress. Untreated pain can affect the animals’ biology and add variability to the experiment; however specific pain management procedures can also introduce variability, affecting experimental data [195, 196]. Under-reporting of welfare management procedures can also contribute to the perpetuation of non-compliant methodologies and insufficient or inappropriate use of analgesia [196] or other welfare measures. A thorough description of the procedures used to alleviate pain, suffering and distress provides practical information for researchers to replicate the method.

Clearly describe pain management strategies, including:

- specific analgesic
- administration method (formulation, route, dose, concentration, volume, frequency, timing, and equipment used)
- rationale for the choice (e.g. animal model, disease/pathology, procedure, mechanism of action, pharmacokinetics, personnel safety)
- protocol modifications to reduce pain, suffering and distress (e.g. changes to the anaesthetic protocol, increased frequency of monitoring, procedural modifications, habituation, etc.)

If analgesics or other welfare measures, reasonably expected for the procedure performed, are not performed for experimental reasons, report the scientific justification [197].

#### Examples

1. *“If piglets developed diarrhea, they were placed on an electrolyte solution and provided supplemental water, and if the diarrhea did not resolve within 48 h, piglets received a single dose of ceftiofur (5.0 mg ceftiofur equivalent/kg of body weight i.m [Excede, Zoetis, Florham Park, NJ]). If fluid loss continued after treatment, piglets then received a single dose of sulfamethoxazole and trimethoprim oral suspension (50 mg/8 mg per mL, Hi-Tech Pharmacal, Amityville, NY) for 3 consecutive days.” [198]*
2. *“One hour before surgery, we administered analgesia to the mice by offering them nut paste (Nutella; Ferrero, Pino Torinese, Italy) containing 1 mg per kg body weight buprenorphine (Temgesic; Schering-Plough Europe, Brussels, Belgium) for voluntary ingestion, as described previously. The mice had been habituated to pure nut paste for 2 d prior to surgery.” [199]*
3. *“If a GCPS score equal or greater than 6 (out of 24) was assigned postoperatively, additional analgesia was provided with methadone 0.1 mg kg-1 IM (or IV if required), and pain reassessed 30 minutes later. The number of methadone doses was recorded.” [44]*

16b. Report any expected or unexpected adverse events.

#### Explanation

Reporting adverse events allows other researchers to minimise the risk of these events occurring in their own studies and to plan appropriate welfare assessments. If the experiment is testing the efficacy of a treatment, the occurrence of adverse events may alter the balance between treatment benefit and risk [33].

Report any adverse events, expected or unexpected, that had a negative impact on the welfare of the animals in the study (e.g. cardiovascular and respiratory depression, CNS disturbance, hypothermia, reduction of food intake).

#### Examples

1. *“Murine lymph node tumors arose in 11 of 12 mice that received N2-transduced human cells. The neo gene could be detected in murine cells as well as in human cells. Significant lymphoproliferation could be seen only in the murine pre-T cells. It took 5 months for murine leukemia to arise; the affected mice displayed symptoms of extreme sickness rapidly, with 5 of the 12 mice becoming moribund on exactly the same day (Figure 3), and 6 others becoming moribund within a 1-month period…Of the 12 mice that had received N2-transduced human cells, 11 had to be killed because they developed visibly enlarged lymph nodes and spleen, hunching, and decrease in body weight, as shown in Figure 3…. The 12th mouse was observed carefully for 14 months; it did not show any signs of leukemia or other adverse events, and had no abnormal tissues when it was autopsied.…The mice were observed at least once daily for signs of illness, which were defined as any one or more of the following: weight loss, hunching, lethargy, rapid breathing, skin discoloration or irregularities, bloating, hemiparesis, visibly enlarged lymph nodes, and visible solid tumors under the skin. Any signs of illness were logged as “adverse events” in the experiment, the mouse was immediately killed, and an autopsy was performed to establish the cause of illness.” [200]*
2. *“Although procedures were based on those reported in the literature, dogs under Protocol 1 displayed high levels of stress and many experienced vomiting. This led us to significantly alter procedures in order to optimize the protocol for the purposes of our own fasting and postprandial metabolic studies.” [201]*

16c. Describe the humane endpoints established for the study and the frequency of monitoring.

#### Explanation

Humane endpoints are predetermined morphological, physiological and/or behavioural signs that define the circumstances under which an animal will be removed from an experimental study. The use of humane endpoints can help minimise harm while allowing the scientific objectives to be achieved [202]. Report the humane endpoints that were established for the specific study, species and strain. Include clear criteria of the clinical signs monitored [127], and clinical signs that led to euthanasia or other defined actions. Include details such as general welfare indicators (e.g. weight loss, reduced food intake, abnormal posture) and procedure-specific welfare indicators (e.g. tumour size in cancer studies [47], sensory motor deficits in stroke studies [203]).

Report the timing and frequency of monitoring, taking into consideration the normal circadian rhythm of the animal and timing of scientific procedures, as well as any increase in the frequency of monitoring (e.g. post-surgery recovery, critical times during disease studies, or following the observation of an adverse event). Publishing score sheets of the clinical signs that were monitored [204] can help guide other researchers to develop clinically relevant welfare assessments, particularly for studies reporting novel procedures.

#### Example

1. *“Both the research team and the veterinary staff monitored animals twice daily. Health was monitored by weight (twice weekly), food and water intake, and general assessment of animal activity, panting, and fur condition…. The maximum size the tumors allowed to grow in the mice before euthanasia was 2000 mm^3^.” [205]*

### Item 17. Interpretation/scientific implications

17a. Interpret the results, taking into account the study objectives and hypotheses, current theory and other relevant studies in the literature.

#### Explanation

It is important to interpret the results of the study in the context of the study objectives (see **item 13 – Objectives**). For hypothesis-testing studies, interpretations should be restricted to the primary outcome (see i**tem 6 – Outcome measures**). Exploratory results derived from additional outcomes should not be described as conclusive, as they may be underpowered and less reliable.

Discuss the findings in the context of current theory, ideally with reference to a relevant systematic review, as individual studies do not provide a complete picture. If a systematic review is not available, take care to avoid selectively citing studies that corroborate the results, or only those that report statistically significant findings [206].

Where appropriate, describe any implications of the experimental methods or research findings for improving welfare standards or reducing the number of animals used in future studies (e.g. the use of a novel approach reduced the results’ variability, thus enabling the use of smaller group sizes without losing statistical power). This may not be the primary focus of the research but reporting this information enables wider dissemination and uptake of refined techniques within the scientific community.

#### Examples

1. *“This is in contrast to data provided by an ‘intra-renal IL-18 overexpression’ model [43], and may reflect an IL-18 concentration exceeding the physiologic range in the latter study.” [207]*
2. *“The new apparatus shows potential for considerably reducing the number of animals used in memory tasks designed to detect potential amnesic properties of new drugs… approximately 43,000 animals have been used in these tasks in the past 5 years but with the application of the continual trials apparatus we estimate that this could have been reduced to 26,000…. with the new paradigm the number of animals needed to obtain reliable results and maintain the statistical power of the tasks is greatly reduced.” [208]*
3. *“In summary, our results show that IL-1Ra protects against brain injury and reduces neuroinflammation when administered peripherally to aged and comorbid animals at reperfusion or 3 hours later. These findings address concerns raised in a recent systematic review on IL-1Ra in stroke [209], and provide further supporting evidence for IL-1Ra as a lead candidate for the treatment of ischemic stroke.” [210]*

17b. Comment on the study limitations including potential sources of bias, limitations of the animal model, and imprecision associated with the results.

#### Explanation

Discussing the limitations of the work is important to place the findings in context, interpret the validity of the results, and ascribe a credibility level to its conclusions [211]. Limitations are unavoidable in scientific research, and describing them is essential to share experience, guide best practice, and aid the design of future experiments.

Discuss the quality of evidence presented in the study, and consider how appropriate the animal model is to the specific research question. A discussion on the rigour of the study design to isolate cause and effect (also known as internal validity [212]) should include whether potential risks of bias have been addressed [9].

#### Examples

1. *“Although in this study we did not sample the source herds, the likelihood of these herds to be Influenza A virus (IAV) positive is high given the commonality of IAV infections in the Midwest. However, we cannot fully rule out the possibility that new gilts became infected with resident viruses after arrival to the herd. Although new gilts were placed into isolated designated areas and procedures were in place to minimize disease transmission (eg. isolation, vaccination), these areas or procedures might not have been able to fully contain infections within the designated areas.” [213]*
2. *“Even though our data demonstrates that sustained systemic TLR9 stimulation aggravates diastolic HF in our model of gene-targeted diastolic HF, there are several limitations as to mechanistic explanations of causality, as well as extrapolations to clinical inflammatory disease states and other HF conditions. First, our pharmacological inflammatory model does not allow discrimination between effects caused by direct cardiac TLR9 stimulation to that of indirect effects mediated by systemic inflammation. Second, although several systemic inflammatory conditions have disturbances in the innate immune system as important features, and some of these again specifically encompassing distorted TLR9 signalling [34], sustained TLR9 stimulation does not necessarily represent a clinically relevant inflammatory condition. Finally, the cardiac myocyte SERCA2a KO model does not adequately represent the molecular basis for, or the clinical features of, diastolic HF.” [214]*

### Item 18. Generalisability/translation

Comment on whether, and how, the findings of this study are likely to generalise to other species or experimental conditions, including any relevance to human biology (where appropriate).

#### Explanation

An important purpose of publishing research findings is to inform future research. In the context of animal studies, this might take the form of further *in vivo* research or another research domain (e.g. human clinical trial). Thoughtful consideration is warranted, as additional unnecessary animal studies are wasteful and unethical. Similarly, human clinical trials initiated based on insufficient or misleading animal research evidence increase research waste and negatively influence the risk-benefit balance for research participants [212, 215].

Consider whether the findings may be used to inform future research in a broadly similar context, or whether enough evidence has been accumulated in the literature to justify further research in another species or in humans. Discuss what (if any) further research may be required to allow generalisation or translation. Discuss and interpret the results in relation to current evidence, and in particular whether similar [216] or otherwise supportive [217] findings have been reported by other groups. Discuss the range of circumstances in which the effect is observed, and factors which may moderate that effect. Such factors could include for example the population (e.g. age, sex, strain, species), the intervention (e.g. different drugs of the same class), and the outcome measured (e.g. different approaches to assessing memory).

#### Examples

1. *“Our results demonstrate that hDBS robustly modulates the mesolimbic network. This finding may hold clinical relevance for hippocampal DBS therapy in epilepsy cases, as connectivity in this network has previously been shown to be suppressed in mTLE. Further research is necessary to investigate potential DBS-induced restoration of MTLE-induced loss of functional connectivity in mesolimbic brain structures.” [218]*
2. *“The tumor suppressor effects of GAS1 had been previously reported in cell cultures or in xenograft models, this is the first work in which the suppressor activity of murine Gas1 is reported for primary tumors in vivo. Recent advances in the design of safe vectors for transgene delivery may result in extrapolating our results to humans and so a promising field of research emerges in the area of hepatic, neoplastic diseases.” [219]*

### Item 19. Protocol registration

Provide a statement indicating whether a protocol (including the research question, key design features, and analysis plan) was prepared before the study, and if and where this protocol was registered.

#### Explanation

Akin to the approach taken for clinical trials, protocol registration has emerged as a mechanism that is likely to improve the transparency of animal research [215, 220, 221]. Registering a protocol before the start of the experiment enables researchers to demonstrate that the hypothesis, approach and analysis were planned in advance and not shaped by data as they emerged; it enhances scientific rigour and protects the researcher against concerns about selective reporting of results [222, 223]. A protocol should consist of a) the question being addressed and the key features of the research that is proposed, such as the hypothesis being tested and the primary outcome measure (if applicable), the statistical analysis plan; and b) the laboratory procedures to be used to perform the planned experiment.

Protocols may be registered with different levels of completeness. For example, in the Registered Report format offered by an increasing number of journals, protocols undergo peer review and if accepted, the journal commits to publishing the completed research regardless of the results obtained [220].

Other online resources include the Open Science Framework [224], which is suitable to deposit PHISPS (Population; Hypothesis; Intervention; Statistical Analysis Plan; Primary; Outcome Measure; Sample Size Calculation) protocols [225] and provide researchers with the flexibility to embargo the preregistration and keep it from public view until the research is published, and selectively share it with reviewers and editors. The EDA can also be used to generate a time-stamped PDF, which sets out key elements of the experimental design [18]. This can be used to demonstrate that the study conduct, analysis and reporting were not unduly driven by emerging data. As a minimum we recommend registering protocols containing all PHISPS components as outlined above.

Provide a statement indicating whether or not any protocol was prepared before the study, and if applicable, the location of its registration. Where there have been deviations from the protocol, describe the rationale for these changes in the publication so that readers can take this into account when assessing the findings.

#### Examples

1. *“A detailed description of all protocols can be found in the Registered Report (Kandela et al., 2015). Additional detailed experimental notes, data, and analysis are available on the Open Science Framework (OSF) (RRID: SCR_003238) (https://osf.io/xu1g2/; Mantis et al., 2016).”[226]*
2. *“To maximise the objectivity of the presented analyses, we preregistered this study with its two hypotheses, its planned methods, and its complete plan of data analysis before the start of data collection (https://osf.io/fh8eq/), and we closely adhered to our plan… All statistical analyses closely followed our preregistered analysis plan (https://osf.io/fh8eq/).” [227]*
3. *“We preregistered our analyses with the Open Science Framework which facilitates reproducibility and open collaboration in science research (Foster & Deardorff, 2017). Our preregistration: Sheldon and Griffith (2017), was carried out to limit the number of analyses conducted and to validate our commitment to testing a limited number of a priori hypotheses. Our methods are consistent with this preregistration (Sheldon & Griffith, 2017).” [228]*

### Item 20. Data access

Provide a statement describing if and where study data are available.

#### Explanation

A data sharing statement describes how others can access the data on which the paper is based. Sharing adequately annotated data allows others to replicate data analyses, so that results can be independently tested and verified. Data sharing allows the data to be repurposed and new datasets to be created by combining data from multiple studies (e.g. to be used in secondary analyses). This allows others to explore new topics and increases the impact of the study, potentially preventing unnecessary use of animals and providing more value for money. Access to raw data also facilitates text and automated data mining [229].

An increasing number of publishers and funding bodies require authors or grant holders to make their data publicly available [230]. Journal articles with accompanying data may be cited more frequently [231]. Datasets can also be independently cited in their own right, which provides additional credit for authors. This practice is gaining increasing recognition and acceptance [232].

Where possible, make available all data that contribute to summary estimates or claims presented in the paper. Data should follow the FAIR guiding principles [233], that is data are findable, accessible (i.e. don’t use outdated file types), interoperable (can be used on multiple platforms and with multiple software packages) and re-usable (i.e. have adequate data descriptors).

Data can be made publicly available via a structured, specialised (domain-specific), open access repository such as those maintained by NCBI (https://www.ncbi.nlm.nih.gov/) or EBI (https://www.ebi.ac.uk/). If such a repository is not available, data can be deposited in unstructured but publicly available repositories (e.g. Figshare (https://figshare.com/), Dryad (https://datadryad.org/), Zenodo (https://zenodo.org/) or Open Science Framework (https://osf.io/)). There are also search platforms to identify relevant repositories with rigorous standards (e.g. FairSharing (https://fairsharing.org/) and re3data (https://www.re3data.org/).

#### Examples

1. *“Data Availability Statement: All data are available from Figshare at http://dx.doi.org/10.6084/m9.figshare.1288935.”[234]*
2. *“A fundamental goal in generating this dataset is to facilitate access to spiny mouse transcript sequence information for external collaborators and researchers. The sequence reads and metadata are available from the NCBI (PRJNA342864) and assembled transcriptomes (Trinity_v2.3.2 and tr2aacds_v2) are available from the Zenodo repository (https://doi.org/10.5281/zenodo.808870), however accessing and utilizing this data can be challenging for researchers lacking bioinformatics expertise. To address this problem we are hosting a SequenceServer32 BLAST-search website (http://spinymouse.erc.monash.edu/sequenceserver/http://spinymouse.erc.monash.edu/sequenceserver/). This resource provides a user-friendly interface to access sequence information from the tr2aacds_v2 assembly (to explore annotated protein-coding transcripts) and/or the Trinity_v2.3.2 assembly (to explore non-coding transcripts).” [235]*

### Item 21. Declaration of interests

21a. Declare any potential conflicts of interest, including financial and non-financial. If none exist, this should be stated.

#### Explanation

A competing or conflict of interest is anything that interferes with (or could be perceived as interfering with) the full and objective presentation, analysis, and interpretation of the research. Competing or conflicts of interest can be financial or non-financial, professional or personal. They can exist in institutions, in teams, or with individuals. Potential competing interests are considered in peer review, editorial and publication decisions; the aim is to ensure transparency, and in most cases, a declaration of a conflict of interest does not obstruct the publication or review process.

Examples are provided in Box 8. If unsure, declare all potential conflicts, including both perceived and real conflicts of interest [236].

##### Box 8: Examples of competing or conflicts of interest

**<u>Financial:</u>**

Funding and other payments received or expected by the authors directly arising from the publication of the study, or funding or other payments from an organisation with an interest in the outcome of the work.

**<u>Non-financial:</u>**

Research that may benefit the individual or institution in terms of goods in kind. This includes unpaid advisory position in a government, non-government organisation or commercial organisations.

**<u>Affiliations:</u>**

Employed by, on the advisory board or a member of an organisation with an interest in the outcome of the work.

**<u>Intellectual property:</u>**

Patents or trademarks owned by someone or their organisation. This also includes the potential exploitation of the scientific advance being reported for the institution, the authors, or the research funders.

**<u>Personal:</u>**

Friends, family, relationships, and other close personal connections to people who may potentially benefit financially or in other ways from the research.

**<u>Ideology:</u>**

Beliefs or activism (e.g. political or religious) relevant to the work. Membership of a relevant advocacy or lobbying organisation.

#### Examples

1. *“The study was funded by Gubra ApS. LSD; PJP; GH; KF and HBH are employed by Gubra ApS. JJ and NV are the owners of Gubra ApS. Gubra ApS provided support in the form of materials and salaries for authors LSD; PJP; GH; KF; HBH; JJ and NV.” [237]*
2. *“The authors have declared that no competing interests exist.” [238]*.

21b. List all funding sources (including grant identifier) and the role of the funder(s) in the design, analysis and reporting of the study.

#### Explanation

The identification of funding sources allows the reader to assess any competing interests, and any potential sources of bias. For example, bias, as indicated by a prevalence of more favourable outcomes, has been demonstrated for clinical research funded by industry compared to studies funded by other sources [239–241]. Evidence for preclinical research also indicates that funding sources may influence the interpretation of study outcomes [236, 242].

Report the funding information including the financial supporting body(s) and any grant identifier(s). Include the role of the funder in the design, analysis, reporting and/or or decision to publish. If the research did not receive specific funding, but was performed as part of the employment of the authors, name the employer.

#### Examples

1. *“Support was provided by the Italian Ministry of Health: Current research funds PRC 2010/001 [http://www.salute.gov.it/] to MG. The funders had no role in study design, data collection and analysis, decision to publish, or preparation of the manuscript.” [243]*
2. *“This study was financially supported by the Tuberculosis and Lung Research Center of Tabriz University of Medical Sciences and the Research Council of University of Tabriz. The funders had no role in study design, data collection and analysis, decision to publish, or preparation of the manuscript.” [244]*
3. *“This work was supported by the salary paid to AEW. The funder had no role in study design, data collection and analysis, decision to publish, or preparation of the manuscript.” [245]*

## Acknowledgements

We would like to acknowledge the late Doug Altman’s contribution to this project, Doug was a dedicated member of the working group and his input into the guidelines’ revision has been invaluable.

## References

1. Kilkenny C, Parsons N, Kadyszewski E, Festing MF, Cuthill IC, Fry D, et al. Survey of the quality of experimental design, statistical analysis and reporting of research using animals. PLoS One. 2009;4(11):e7824. Epub 2009/12/04. doi: 10.1371/journal.pone.0007824. PubMed PMID: 19956596; PubMed Central PMCID: PMCPMC2779358.

2. Florez-Vargas O, Brass A, Karystianis G, Bramhall M, Stevens R, Cruickshank S, et al. Bias in the reporting of sex and age in biomedical research on mouse models. Elife. 2016;5. Epub 2016/03/05. doi: 10.7554/eLife.13615. PubMed PMID: 26939790; PubMed Central PMCID: PMCPMC4821800.

3. Weissgerber TL, Garcia-Valencia O, Garovic VD, Milic NM, Winham SJ. Why we need to report more than ‘data were analyzed by t-tests or anova’. Elife. 2018;7. Epub 2018/12/24. doi: 10.7554/eLife.36163. PubMed PMID: 30574870; PubMed Central PMCID: PMCPMC6326723.

4. Macleod MR, Lawson McLean A, Kyriakopoulou A, Serghiou S, de Wilde A, Sherratt N, et al. Risk of bias in reports of in vivo research: A focus for improvement. PLoS Biol. 2015;13(10):e1002273. Epub 2015/10/16. doi: 10.1371/journal.pbio.1002273. PubMed PMID: 26460723; PubMed Central PMCID: PMCPMC4603955.

5. Kilkenny C, Browne WJ, Cuthill IC, Emerson M, Altman DG. Improving bioscience research reporting: The ARRIVE guidelines for reporting animal research. PLoS Biol. 2010;8(6):e1000412. Epub 2010/07/09. doi: 10.1371/journal.pbio.1000412. PubMed PMID: 20613859; PubMed Central PMCID: PMCPMC2893951.

6. Baker D, Lidster K, Sottomayor A, Amor S. Two years later: Journals are not yet enforcing the ARRIVE guidelines on reporting standards for pre-clinical animal studies. PLoS Biol. 2014;12(1):e1001756. Epub 2014/01/11. doi: 10.1371/journal.pbio.1001756. PubMed PMID: 24409096; PubMed Central PMCID: PMCPMC3883646.

7. Gulin JE, Rocco DM, Garcia-Bournissen F. Quality of reporting and adherence to ARRIVE guidelines in animal studies for chagas disease preclinical drug research: A systematic review. PLoS Negl Trop Dis. 2015;9(11):e0004194. Epub 2015/11/21. doi: 10.1371/journal.pntd.0004194. PubMed PMID: 26587586; PubMed Central PMCID: PMCPMC4654562.

8. Avey MT, Moher D, Sullivan KJ, Fergusson D, Griffin G, Grimshaw JM, et al. The devil is in the details: Incomplete reporting in preclinical animal research. PLoS ONE. 2016;11(11):e0166733. doi: 10.1371/journal.pone.0166733.

9. Reichlin TS, Vogt L, Würbel H. The researchers’ view of scientific rigor—survey on the conduct and reporting of in vivo research. PLoS ONE. 2016;11(12):e0165999. doi: 10.1371/journal.pone.0165999.

10. Leung V, Rousseau-Blass F, Beauchamp G, Pang DSJ. ARRIVE has not ARRIVEd: Support for the ARRIVE (animal research: Reporting of in vivo experiments) guidelines does not improve the reporting quality of papers in animal welfare, analgesia or anesthesia. PLoS One. 2018;13(5):e0197882. Epub 2018/05/26. doi: 10.1371/journal.pone.0197882. PubMed PMID: 29795636; PubMed Central PMCID: PMCPMC5967836.

11. Hair K, Macleod MR, Sena ES, Sena ES, Hair K, Macleod MR, et al. A randomised controlled trial of an intervention to improve compliance with the ARRIVE guidelines (iicarus). Research Integrity and Peer Review. 2019;4(1):12. doi: 10.1186/s41073-019-0069-3.

12. McGrath JC, Lilley E. Implementing guidelines on reporting research using animals (ARRIVE etc.): New requirements for publication in bjp. Br J Pharmacol. 2015;172(13):3189–93. Epub 2015/05/13. doi: 10.1111/bph.12955. PubMed PMID: 25964986; PubMed Central PMCID: PMCPMC4500358.

13. Percie du Sert N, Hurst V, Ahluwalia A, Alam S, Avey MT, Baker M, et al. The ARRIVE guidelines 2019: Updated guidelines for reporting animal research. Submitted to Plos Biology. 2019.

14. van der Worp HB, Howells DW, Sena ES, Porritt MJ, Rewell S, O’Collins V, et al. Can animal models of disease reliably inform human studies? PLOS Medicine. 2010;7(3):e1000245. doi: 10.1371/journal.pmed.1000245.

15. NC3Rs. Experimental design assistant version 1 [22/05/2019]. Available from: https://eda.nc3rs.org.uk/.

16. Ruxton G. Experimental design for the life sciences. New York, NY: Oxford University Press; 2016. pages cm p.

17. Festing MF, Altman DG. Guidelines for the design and statistical analysis of experiments using laboratory animals. ILAR J. 2002;43(4):244–58. Epub 2002/10/23. PubMed PMID: 12391400.

18. Percie du Sert N, Bamsey I, Bate ST, Berdoy M, Clark RA, Cuthill I, et al. The experimental design assistant. PLoS Biol. 2017;15(9):e2003779. Epub 2017/09/29. doi: 10.1371/journal.pbio.2003779. PubMed PMID: 28957312; PubMed Central PMCID: PMCPMC5634641.

19. Nguyen TT, Bazzoli C, Mentre F. Design evaluation and optimisation in crossover pharmacokinetic studies analysed by nonlinear mixed effects models. Stat Med. 2012;31(11-12):1043–58. Epub 2011/10/04. doi: 10.1002/sim.4390. PubMed PMID: 21965170.

20. Hill D, Chen L, Snaar-Jagalska E, Chaudhry B. Embryonic zebrafish xenograft assay of human cancer metastasis [version 2; referees: 2 approved]. F1000Research. 2018;7(1682). doi: 10.12688/f1000research.16659.2.

21. Hurlbert SH. Pseudoreplication and the design of ecological field experiments. Ecological Monographs. 1984;54(2):187–211. doi: 10.2307/1942661.

22. Lazic SE. The problem of pseudoreplication in neuroscientific studies: Is it affecting your analysis? BMC Neuroscience. 2010;11(1):5. doi: 10.1186/1471-2202-11-5.

23. Burdge GC, Lillycrop KA, Jackson AA, Gluckman PD, Hanson MA. The nature of the growth pattern and of the metabolic response to fasting in the rat are dependent upon the dietary protein and folic acid intakes of their pregnant dams and post-weaning fat consumption. Br J Nutr. 2008;99(3):540–9. Epub 2007/09/01. doi: 10.1017/S0007114507815819. PubMed PMID: 17761015; PubMed Central PMCID: PMCPMC2493056.

24. Bate ST, Clark RA. The design and statistical analysis of animal experiments. Cambridge, United Kingdom: Cambridge University Press; 2014. xv, 310 pages p.

25. Lazic SE, Clarke-Williams CJ, Munafò MR. What exactly is ‘n’ in cell culture and animal experiments? PLOS Biology. 2018;16(4):e2005282. doi: 10.1371/journal.pbio.2005282.

26. NC3Rs. Experimental unit 2015 [cited 2019 21 March]. Available from: https://eda.nc3rs.org.uk/experimental-design-unit.

27. Kwan S, King J, Grenier J, Yan J, Jiang X, Roberson M, et al. Maternal choline supplementation during normal murine pregnancy alters the placental epigenome: Results of an exploratory study. Nutrients. 2018;10(4):417. PubMed PMID: doi:10.3390/nu10040417.

28. Karp NA, Mason J, Beaudet AL, Benjamini Y, Bower L, Braun RE, et al. Prevalence of sexual dimorphism in mammalian phenotypic traits. Nature Communications. 2017;8:15475. doi: 10.1038/ncomms15475.

29. Ribeiro FdAS, Vasquez LA, Fernandes JBK, Sakomura NK. Feeding level and frequency for freshwater angelfish. Revista Brasileira de Zootecnia. 2012;41:1550–4.

30. Grasselli G, Rossi S, Musella A, Gentile A, Loizzo S, Muzio L, et al. Abnormal nmda receptor function exacerbates experimental autoimmune encephalomyelitis. British journal of pharmacology. 2013;168(2):502–17. doi: 10.1111/j.1476-5381.2012.02178.x. PubMed PMID: 22924679.

31. Gray LE, Jr., Ostby J, Furr J, Price M, Veeramachaneni DN, Parks L. Perinatal exposure to the phthalates dehp, bbp, and dinp, but not dep, dmp, or dotp, alters sexual differentiation of the male rat. Toxicol Sci. 2000;58(2):350–65. Epub 2000/12/02. PubMed PMID: 11099647.

32. Vahidy F, Schäbitz W-R, Fisher M, Aronowski J. Reporting standards for preclinical studies of stroke therapy. Stroke. 2016;47(10):2435.

33. Muhlhausler BS, Bloomfield FH, Gillman MW. Whole animal experiments should be more like human randomized controlled trials. PLoS Biol. 2013;11(2):e1001481. doi: 10.1371/journal.pbio.1001481.

34. Jennions MD, Møller AP. A survey of the statistical power of research in behavioral ecology and animal behavior. Behavioral Ecology. 2003;14(3):438–45. doi: 10.1093/beheco/14.3.438.

35. Button KS, Ioannidis JPA, Mokrysz C, Nosek BA, Flint J, Robinson ESJ, et al. Power failure: Why small sample size undermines the reliability of neuroscience. Nat Rev Neurosci. 2013;14(5):365–76. doi: 10.1038/nrn3475.

36. Würbel H. More than 3rs: The importance of scientific validity for harm-benefit analysis of animal research. Lab animal. 2017;46(4):164.

37. Walker M, Fureix C, Palme R, Newman JA, Ahloy Dallaire J, Mason G. Mixed-strain housing for female c57bl/6, dba/2, and balb/c mice: Validating a split-plot design that promotes refinement and reduction. BMC Medical Research Methodology. 2016;16(1):11. doi: 10.1186/s12874-016-0113-7.

38. Shaw R, Festing MF, Peers I, Furlong L. Use of factorial designs to optimize animal experiments and reduce animal use. ILAR J. 2002;43(4):223–32. Epub 2002/10/23. PubMed PMID: 12391398.

39. Neumann K, Grittner U, Piper SK, Rex A, Florez-Vargas O, Karystianis G, et al. Increasing efficiency of preclinical research by group sequential designs. PLoS Biol. 2017;15(3):e2001307. Epub 2017/03/11. doi: 10.1371/journal.pbio.2001307. PubMed PMID: 28282371; PubMed Central PMCID: PMCPMC5345756.

40. Faul F, Erdfelder E, Lang AG, Buchner A. G*power 3: A flexible statistical power analysis program for the social, behavioral, and biomedical sciences. Behav Res Methods. 2007;39(2):175–91. Epub 2007/08/19. PubMed PMID: 17695343.

41. Festing MFW. On determining sample size in experiments involving laboratory animals. Laboratory Animals. 2018;52(4):341–50. doi: 10.1177/0023677217738268.

42. Reynolds PS, McCarter J, Sweeney C, Mohammed BM, Brophy DF, Fisher B, et al. Informing efficient pilot development of animal trauma models through quality improvement strategies. Lab Anim. 2018:23677218802999. Epub 2018/10/10. doi: 10.1177/0023677218802999. PubMed PMID: 30296892.

43. Bate S. How to decide your sample size when the power calculation is not straightforward NC3Rs.org.uk: NC3Rs; 2018 [updated 01/08/2018; cited 2018 02/08/2018]. Available from: https://www.nc3rs.org.uk/news/how-decide-your-sample-size-when-power-calculation-not-straightforward.

44. Bustamante R, Daza MA, Canfrán S, García P, Suárez M, Trobo I, et al. Comparison of the postoperative analgesic effects of cimicoxib, buprenorphine and their combination in healthy dogs undergoing ovariohysterectomy. Veterinary Anaesthesia and Analgesia. 2018;45(4):545–56. doi: 10.1016/j.vaa.2018.01.003.

45. Spin JR, Oliveira GJPLd, Spin-Neto R, Pires JR, Tavares HS, Ykeda F, et al. Avaliação histomorfométrica da associação entre biovidro e osso bovino liofilizado no tratamento de defeitos ósseos críticos criados em calvárias de ratos. Estudo piloto. Revista de Odontologia da UNESP. 2015;44:37–43.

46. Salkind NJ. Encyclopedia of research design: Sage; 2010.

47. Workman P, Aboagye EO, Balkwill F, Balmain A, Bruder G, Chaplin DJ, et al. Guidelines for the welfare and use of animals in cancer research. British Journal Of Cancer. 2010;102:1555. doi: 10.1038/sj.bjc.6605642.

48. Sena ES, Jeffreys AL, Cox SF, Sastra SA, Churilov L, Rewell S, et al. The benefit of hypothermia in experimental ischemic stroke is not affected by pethidine. International journal of stroke: official journal of the International Stroke Society. 2013;8(3):180–5. doi: 10.1111/j.1747-4949.2012.00834.x. PubMed PMID: 22759525.

49. Kafkafi N, Agassi J, Chesler EJ, Crabbe JC, Crusio WE, Eilam D, et al. Reproducibility and replicability of rodent phenotyping in preclinical studies. Neuroscience & Biobehavioral Reviews. 2018;87:218–32. doi: 10.1016/j.neubiorev.2018.01.003.

50. Scott S, Kranz JE, Cole J, Lincecum JM, Thompson K, Kelly N, et al. Design, power, and interpretation of studies in the standard murine model of als. Amyotrophic Lateral Sclerosis. 2008;9(1):4–1 5. doi: 10.1080/17482960701856300.

51. Holman C, Piper SK, Grittner U, Diamantaras AA, Kimmelman J, Siegerink B, et al. Where have all the rodents gone? The effects of attrition in experimental research on cancer and stroke. PLoS Biol. 2016;14(1):e1002331. Epub 2016/01/05. doi: 10.1371/journal.pbio.1002331. PubMed PMID: 26726833; PubMed Central PMCID: PMCPMC4699644.

52. Rice ASC, Morland R, Huang W, Currie GL, Sena ES, Macleod MR. Transparency in the reporting of in vivo pre-clinical pain research: The relevance and implications of the ARRIVE (animal research: Reporting in vivo experiments) guidelines. Scandinavian Journal of Pain. 2013;4(2):58–62. doi: 10.1016/j.sjpain.2013.02.002.

53. Genther-Schroeder ON, Branine ME, Hansen SL. Effects of increasing supplemental dietary zn concentration on growth performance and carcass characteristics in finishing steers fed ractopamine hydrochloride. J Anim Sci. 2018;96(5):1903–13. Epub 2018/05/08. doi: 10.1093/jas/sky094. PubMed PMID: 29733414.

54. Castiglioni L, Colazzo F, Fontana L, Colombo GI, Piacentini L, Bono E, et al. Evaluation of left ventricle function by regional fractional area change (rfac) in a mouse model of myocardial infarction secondary to valsartan treatment. PLOS ONE. 2015;10(8):e0135778. doi: 10.1371/journal.pone.0135778.

55. Brent LJ, Heilbronner SR, Horvath JE, Gonzalez-Martinez J, Ruiz-Lambides A, Robinson AG, et al. Genetic origins of social networks in rhesus macaques. Sci Rep. 2013;3:1042. Epub 2013/01/11. doi: 10.1038/srep01042. PubMed PMID: 23304433; PubMed Central PMCID: PMCPMC3540398.

56. Gouveia K, Hurst JL. Optimising reliability of mouse performance in behavioural testing: The major role of non-aversive handling. Sci Rep. 2017;7:44999. Epub 2017/03/23. doi: 10.1038/srep44999. PubMed PMID: 28322308; PubMed Central PMCID: PMC5359560.

57. Schulz KF, Chalmers I, Hayes RJ, Altman DG. Empirical evidence of bias. Dimensions of methodological quality associated with estimates of treatment effects in controlled trials. Jama. 1995;273(5):408–12. Epub 1995/02/01. PubMed PMID: 7823387.

58. Schulz KF, Grimes DA. Allocation concealment in randomised trials: Defending against deciphering. Lancet. 2002;359(9306):614–8. Epub 2002/02/28. doi: 10.1016/s0140-6736(02)07750-4. PubMed PMID: 11867132.

59. Chalmers TC, Celano P, Sacks HS, Smith Jr H. Bias in treatment assignment in controlled clinical trials. New England Journal of Medicine. 1983;309(22):1 358–61.

60. Hirst JA, Howick J, Aronson JK, Roberts N, Perera R, Koshiaris C, et al. The need for randomization in animal trials: An overview of systematic reviews. PLoS ONE. 2014;9(6):e98856. doi: 10.1371/journal.pone.0098856.

61. Vesterinen HM, Sena ES, ffrench-Constant C, Williams A, Chandran S, Macleod MR. Improving the translational hit of experimental treatments in multiple sclerosis. Multiple Sclerosis Journal. 2010;16(9):1044–55. doi: doi:10.1177/1352458510379612. PubMed PMID: 20685763.

62. Taves DR. Minimization: A new method of assigning patients to treatment and control groups. Clin Pharmacol Ther. 1974;15(5):443–53. Epub 1974/05/01. PubMed PMID: 4597226.

63. Saint-Mont U. Randomization does not help much, comparability does. PLOS ONE. 2015;10(7):e0132102. doi: 10.1371/journal.pone.0132102.

64. Laajala TD, Jumppanen M, Huhtaniemi R, Fey V, Kaur A, Knuuttila M, et al. Optimized design and analysis of preclinical intervention studies in vivo. Scientific Reports. 2016;6:30723. doi: 10.1038/srep30723.

65. Zhao S, Kang R, Deng T, Luo L, Wang J, Li E, et al. Comparison of two cannulation methods for assessment of intracavernosal pressure in a rat model. PLoS One. 2018;13(2):e0193543. Epub 2018/02/28. doi: 10.1371/journal.pone.0193543. PubMed PMID: 29486011; PubMed Central PMCID: PMCPMC5828359.

66. Jansen of Lorkeers SJ, Gho JMIH, Koudstaal S, van Hout GPJ, Zwetsloot PPM, van Oorschot JWM, et al. Xenotransplantation of human cardiomyocyte progenitor cells does not improve cardiac function in a porcine model of chronic ischemic heart failure. Results from a randomized, blinded, placebo controlled trial. PLOS ONE. 2015;10(12):e0143953. doi: 10.1371/journal.pone.0143953.

67. Ishida A, Mutoh T, Ueyama T, Bando H, Masubuchi S, Nakahara D, et al. Light activates the adrenal gland: Timing of gene expression and glucocorticoid release. Cell Metab. 2005;2(5):297–307. Epub 2005/11/08. doi: 10.1016/j.cmet.2005.09.009. PubMed PMID: 16271530.

68. Kang M, Ragan BG, Park JH. Issues in outcomes research: An overview of randomization techniques for clinical trials. J Athl Train. 2008;43(2):215–21. Epub 2008/03/18. doi: 10.4085/1062-6050-43.2.215. PubMed PMID: 18345348; PubMed Central PMCID: PMCPMC2267325.

69. Altman DG, Bland JM. Statistics notes. Treatment allocation in controlled trials: Why randomise? BMJ. 1999;318(7192):1209. Epub 1999/04/30. PubMed PMID: 10221955; PubMed Central PMCID: PMCPMC1115595.

70. Altman DG, Bland JM. Treatment allocation by minimisation. BMJ. 2005;330(7495):843. Epub 2005/04/09. doi: 10.1136/bmj.330.7495.843. PubMed PMID: 15817555; PubMed Central PMCID: PMCPMC556084.

71. El-Agroudy NN, El-Naga RN, El-Razeq RA, El-Demerdash E. Forskolin, a hedgehog signalling inhibitor, attenuates carbon tetrachloride-induced liver fibrosis in rats. British Journal of Pharmacology. 2016;173(22):3248–60. doi: 10.1111/bph.13611.

72. Carrillo M, Migliorati F, Bruls R, Han Y, Heinemans M, Pruis I, et al. Repeated witnessing of conspecifics in pain: Effects on emotional contagion. PLOS ONE. 2015;10(9):e0136979. doi: 10.1371/journal.pone.0136979.

73. Del Bianco Benedeti P, Paulino PVR, Marcondes MI, Maciel IFS, da Silva MC, Faciola AP. Partial replacement of ground corn with glycerol in beef cattle diets: Intake, digestibility, performance, and carcass characteristics. PLOS ONE. 2016;11(1):e0148224. doi: 10.1371/journal.pone.0148224.

74. Nuzzo R. How scientists fool themselves–and how they can stop. Nature News. 2015;526(7572):182.

75. Hróbjartsson A, Thomsen ASS, Emanuelsson F, Tendal B, Hilden J, Boutron I, et al. Observer bias in randomised clinical trials with binary outcomes: Systematic review of trials with both blinded and non-blinded outcome assessors. BMJ. 2012;344. doi: 10.1136/bmj.e1119.

76. Rosenthal R, Fode KL. The effect of experimenter bias on the performance of the albino rat. Behavioral Science. 1963;8(3):183–9. doi: 10.1002/bs.3830080302.

77. Rosenthal R, Lawson R. A longitudinal study of the effects of experimenter bias on the operant learning of laboratory rats. Journal of Psychiatric Research. 1964;2(2):61–72. doi: 10.1016/0022-3956(64)90003-2.

78. Macleod MR, van der Worp HB, Sena ES, Howells DW, Dirnagl U, Donnan GA. Evidence for the efficacy of nxy-059 in experimental focal cerebral ischaemia is confounded by study quality. Stroke. 2008;39(10):2824–9. Epub 2008/07/19. doi: 10.1161/strokeaha.108.515957. PubMed PMID: 18635842.

79. Vesterinen HM, Sena ES, ffrench-Constant C, Williams A, Chandran S, Macleod MR. Improving the translational hit of experimental treatments in multiple sclerosis. Mult Scler. 2010;16(9):1044–55. Epub 2010/08/06. doi: 1352458510379612 [pii] 10.1177/1352458510379612. PubMed PMID: 20685763.

80. Gómez de Segura IA, Bustamante R, Daza MA, Canfrán S, García P, Suárez M, et al. Comparison of the postoperative analgesic effects of cimicoxib, buprenorphine and their combination in healthy dogs undergoing ovariohysterectomy. Veterinary Anaesthesia and Analgesia. 2018. doi: 10.1016/j.vaa.2018.01.003.

81. Neumann A-M, Abele J, Kirschstein T, Engelmann R, Sellmann T, Köhling R, et al. Mycophenolate mofetil prevents the delayed t cell response after pilocarpine-induced status epilepticus in mice. PLOS ONE. 2017;12(11):e0187330. doi: 10.1371/journal.pone.0187330.

82. Hsieh LS, Wen JH, Miyares L, Lombroso PJ, Bordey A. Outbred cd1 mice are as suitable as inbred c57bl/6j mice in performing social tasks. Neuroscience Letters. 2017;637:142–7. doi: 10.1016/j.neulet.2016.11.035.

83. Claudino MA, Leiria LOS, da Silva FH, Alexandre EC, Renno A, Mónica FZ, et al. Urinary bladder dysfunction in transgenic sickle cell disease mice. PLOS ONE. 2015;10(8):e0133996. doi: 10.1371/journal.pone.0133996.

84. John LK, Loewenstein G, Prelec D. Measuring the prevalence of questionable research practices with incentives for truth telling. Psychological Science. 2012;23(5):524–32. doi: 10.1177/0956797611430953.

85. Landis SC, Amara SG, Asadullah K, Austin CP, Blumenstein R, Bradley EW, et al. A call for transparent reporting to optimize the predictive value of preclinical research. Nature. 2012;490(7419):187–91. Epub 2012/10/13. doi: 10.1038/nature11556. PubMed PMID: 23060188; PubMed Central PMCID: PMCPMC3511845.

86. Head ML, Holman L, Lanfear R, Kahn AT, Jennions MD. The extent and consequences of p-hacking in science. PLOS Biology. 2015;13(3):e1002106. doi: 10.1371/journal.pbio.1002106.

87. Munafò MR, Nosek BA, Bishop DVM, Button KS, Chambers CD, Percie du Sert N, et al. A manifesto for reproducible science. Nature Human Behaviour. 2017;1:0021. doi: 10.1038/s41562-016-0021.

88. Emmrich J, Neher J, Boehm-Sturm P, Endres M, Dirnagl U, Harms C. Stage 1 registered report: Effect of deficient phagocytosis on neuronal survival and neurological outcome after temporary middle cerebral artery occlusion (tmcao) [version 3; referees: 2 approved]. F1000Research. 2018;6(1827). doi: 10.12688/f1000research.12537.3.

89. Lang TA, Altman DG. Basic statistical reporting for articles published in biomedical journals: The “statistical analyses and methods in the published literature” or the sampl guidelines. Int J Nurs Stud. 2015;52(1):5–9. Epub 2014/12/03. doi: 10.1016/j.ijnurstu.2014.09.006. PubMed PMID: 25441757.

90. Polo J, Rodríguez C, Ródenas J, Russell LE, Campbell JM, Crenshaw JD, et al. Ultraviolet light (uv) inactivation of porcine parvovirus in liquid plasma and effect of uv irradiated spray dried porcine plasma on performance of weaned pigs. PLOS ONE. 2015;10(7):e0133008. doi: 10.1371/journal.pone.0133008.

91. Garner JP, Dufour B, Gregg LE, Weisker SM, Mench JA. Social and husbandry factors affecting the prevalence and severity of barbering (‘whisker trimming’) by laboratory mice. Applied Animal Behaviour Science. 2004;89(3-4):263–82. doi: 10.1016/j.applanim.2004.07.004. PubMed PMID: WOS:000225168300007.

92. Podrini C, Cambridge EL, Lelliott CJ, Carragher DM, Estabel J, Gerdin A-K, et al. High-fat feeding rapidly induces obesity and lipid derangements in c57bl/6n mice. Mammalian genome: official journal of the International Mammalian Genome Society. 2013;24(5-6):240–51. doi: 10.1007/s00335-013-9456-0. PubMed PMID: 23712496.

93. Ohishi T, Wang L, Akane H, Shiraki A, Itahashi M, Mitsumori K, et al. Transient suppression of late-stage neuronal progenitor cell differentiation in the hippocampal dentate gyrus of rat offspring after maternal exposure to nicotine. Arch Toxicol. 2014;88(2):443–54. Epub 2013/07/31. doi: 10.1007/s00204-013-1100-y. PubMed PMID: 23892646.

94. Ruxton G, Colegrave N. Experimental design for the life sciences. Fourth ed: Oxford University Press; 2017.

95. Nemeth M, Millesi E, Wagner K-H, Wallner B. Sex-specific effects of diets high in unsaturated fatty acids on spatial learning and memory in guinea pigs. PLOS ONE. 2015;10(10):e0140485. doi: 10.1371/journal.pone.0140485.

96. Keck VA, Edgerton DS, Hajizadeh S, Swift LL, Dupont WD, Lawrence C, et al. Effects of habitat complexity on pair-housed zebrafish. J Am Assoc Lab Anim Sci. 2015;54(4):378–83. Epub 2015/08/01. PubMed PMID: 26224437; PubMed Central PMCID: PMCPMC4521571.

97. Clayton JA, Collins FS. Policy: Nih to balance sex in cell and animal studies. Nature News. 2014;509(7500):282.

98. Shapira S, Sapir M, Wengier A, Grauer E, Kadar T. Aging has a complex effect on a rat model of ischemic stroke. Brain research. 2002;925(2):148–58. doi: 10.1016/s0006-8993(01)03270-x. PubMed PMID: 11792363.

99. Vital M, Harkema JR, Rizzo M, Tiedje J, Brandenberger C. Alterations of the murine gut microbiome with age and allergic airway disease. J Immunol Res. 2015;2015:892568. Epub 2015/06/20. doi: 10.1155/2015/892568. PubMed PMID: 26090504; PubMed Central PMCID: PMCPMC4451525.

100. Bouwknecht JA, Paylor R. Behavioral and physiological mouse assays for anxiety: A survey in nine mouse strains. Behav Brain Res. 2002;136(2):489–501. Epub 2002/11/14. PubMed PMID: 12429412.

101. Simon MM, Greenaway S, White JK, Fuchs H, Gailus-Durner V, Wells S, et al. A comparative phenotypic and genomic analysis of c57bl/6j and c57bl/6n mouse strains. Genome Biol. 2013;14(7):R82. Epub 2013/08/02. doi: 10.1186/gb-2013-14-7-r82. PubMed PMID: 23902802; PubMed Central PMCID: PMCPMC4053787.

102. Jackson SJ, Andrews N, Ball D, Bellantuono I, Gray J, Hachoumi L, et al. Does age matter? The impact of rodent age on study outcomes. Laboratory Animals. 2017;51(2):160–9. doi: 10.1177/0023677216653984. PubMed PMID: 27307423.

103. Khokha MK, Chung C, Bustamante EL, Gaw LW, Trott KA, Yeh J, et al. Techniques and probes for the study of xenopus tropicalis development. Dev Dyn. 2002;225(4):499–510. Epub 2002/11/28. doi: 10.1002/dvdy.10184. PubMed PMID: 12454926.

104. Okva K, Nevalainen T, Pokk P. The effect of cage shelf on the behaviour of male c57bl/6 and balb/c mice in the elevated plus maze test. Lab Anim. 2013;47(3):220–2. Epub 2013/06/14. doi: 10.1177/0023677213489280. PubMed PMID: 23760964.

105. Akkers RC, van Heeringen SJ, Jacobi UG, Janssen-Megens EM, Francoijs KJ, Stunnenberg HG, et al. A hierarchy of h3k4me3 and h3k27me3 acquisition in spatial gene regulation in xenopus embryos. Dev Cell. 2009;17(3):425–34. Epub 2009/09/18. doi: 10.1016/j.devcel.2009.08.005. PubMed PMID: 19758566; PubMed Central PMCID: PMCPMC2746918.

106. Mahler Convenor M, Berard M, Feinstein R, Gallagher A, Illgen-Wilcke B, Pritchett-Corning K, et al. Felasa recommendations for the health monitoring of mouse, rat, hamster, guinea pig and rabbit colonies in breeding and experimental units. Lab Anim. 2014;48(3):178–92. Epub 2014/02/06. doi: 10.1177/0023677213516312. PubMed PMID: 24496575.

107. Baker DG. Natural pathogens of laboratory mice, rats, and rabbits and their effects on research. Clinical Microbiology Reviews. 1998;11(2):231–66. doi: 10.1128/cmr.11.2.231.

108. Velazquez EM, Nguyen H, Heasley KT, Saechao CH, Gil LM, Rogers AWL, et al. Endogenous enterobacteriaceae underlie variation in susceptibility to salmonella infection. Nat Microbiol. 2019. Epub 2019/03/27. doi: 10.1038/s41564-019-0407-8. PubMed PMID: 30911125.

109. Holmdahl R, Malissen B. The need for littermate controls. Eur J Immunol. 2012;42(1):45–7. Epub 2012/01/04. doi: 10.1002/eji.201142048. PubMed PMID: 22213045.

110. Mallapaty S. In the name of reproducibility. Lab Animal. 2018;47(7):178–81. doi: 10.1038/s41684-018-0095-7.

111. Sundberg JP, Schofield PN. Commentary: Mouse genetic nomenclature:Standardization of strain, gene, and protein symbols. Veterinary Pathology. 2010;47(6):1100–4. doi: 10.1177/0300985810374837. PubMed PMID: 20685919.

112. Montoliu L, Whitelaw CBA. Using standard nomenclature to adequately name transgenes, knockout gene alleles and any mutation associated to a genetically modified mouse strain. Transgenic Research. 2011;20(2):435–40. doi: 10.1007/s11248-010-9428-z.

113. Huang W, Feng Y, Liang J, Yu H, Wang C, Wang B, et al. Loss of microrna-128 promotes cardiomyocyte proliferation and heart regeneration. Nat Commun. 2018;9(1):700. Epub 2018/02/18. doi: 10.1038/s41467-018-03019-z. PubMed PMID: 29453456; PubMed Central PMCID: PMCPMC5816015.

114. Ranson A, Cheetham CEJ, Fox K, Sengpiel F. Homeostatic plasticity mechanisms are required for juvenile, but not adult, ocular dominance plasticity. Proceedings of the National Academy of Sciences. 2012;109(4):1311–6. doi: 10.1073/pnas.1112204109.

115. Clarkson JM, Dwyer DM, Flecknell PA, Leach MC, Rowe C. Handling method alters the hedonic value of reward in laboratory mice. Sci Rep. 2018;8(1):2448. Epub 2018/02/07. doi: 10.1038/s41598-018-20716-3. PubMed PMID: 29402923; PubMed Central PMCID: PMCPMC5799408.

116. Hurst JL, West RS. Taming anxiety in laboratory mice. Nat Methods. 2010;7(10):825–6. Epub 2010/09/14. doi: 10.1038/nmeth.1500. PubMed PMID: 20835246.

117. Hewitt JA, Brown LL, Murphy SJ, Grieder F, Silberberg SD. Accelerating biomedical discoveries through rigor and transparency. ILAR J. 2017;58(1):115–28. Epub 2017/06/03. doi: 10.1093/ilar/ilx011. PubMed PMID: 28575443; PubMed Central PMCID: PMCPMC6279133.

118. Almeida JL, Cole KD, Plant AL. Standards for cell line authentication and beyond. PLoS Biol. 2016;14(6):e1002476. Epub 2016/06/15. doi: 10.1371/journal.pbio.1002476. PubMed PMID: 27300367; PubMed Central PMCID: PMCPMC4907466.

119. Leary SL, Underwood W, Anthony R, Cartner S, Corey D, Grandin T, et al. Avma guidelines for the euthanasia of animals: 2013 edition. 2013.

120. Bandrowski AE, Martone ME. Rrids: A simple step toward improving reproducibility through rigor and transparency of experimental methods. Neuron. 2016;90(3):434–6. Epub 2016/05/07. doi: 10.1016/j.neuron.2016.04.030. PubMed PMID: 27151636; PubMed Central PMCID: PMCPMC5854161.

121. Bandrowski A, Brush M, Grethe JS, Haendel MA, Kennedy DN, Hill S, et al. The resource identification initiative: A cultural shift in publishing. J Comp Neurol. 2016;524(1):8–22. Epub 2015/11/26. doi: 10.1002/cne.23913. PubMed PMID: 26599696; PubMed Central PMCID: PMCPMC4684178.

122. Teytelman L, Stoliartchouk A. Protocols.Io: Reducing the knowledge that perishes because we do not publish it. Information Services & Use. 2016;35(1/2):109–15. PubMed PMID: 109101228.

123. Reynolds PS, Fisher BJ, McCarter J, Sweeney C, Martin EJ, Middleton P, et al. Interventional vitamin c: A strategy for attenuation of coagulopathy and inflammation in a swine polytrauma model. J Trauma Acute Care Surg. 2018. Epub 2018/03/15. doi: 10.1097/ta.0000000000001844. PubMed PMID: 29538225.

124. Bauters D, Bedossa P, Lijnen HR, Hemmeryckx B. Functional role of adamts5 in adiposity and metabolic health. PLoS One. 2018;13(1):e0190595. Epub 2018/01/03. doi: 10.1371/journal.pone.0190595. PubMed PMID: 29293679; PubMed Central PMCID: PMCPMC5749841.

125. Bartlang MS, Neumann ID, Slattery DA, Uschold-Schmidt N, Kraus D, Helfrich-Förster C, et al. Time matters: Pathological effects of repeated psychosocial stress during the active, but not inactive, phase of male mice. Journal of Endocrinology. 2012;215(3):425–37. doi: 10.1530/joe-12-0267.

126. Paul AK, Gueven N, Dietis N. Morphine dosing strategy plays a key role in the generation and duration of the produced antinociceptive tolerance. Neuropharmacology. 2017;121:158–66. Epub 2017/04/30. doi: 10.1016/j.neuropharm.2017.04.034. PubMed PMID: 28450061.

127. Hawkins P, Morton DB, Burman O, Dennison N, Honess P, Jennings M, et al. A guide to defining and implementing protocols for the welfare assessment of laboratory animals: Eleventh report of the bvaawf/frame/rspca/ufaw joint working group on refinement. Laboratory Animals. 2011;45(1):1–13. doi: 10.1258/la.2010.010031.

128. Hagemo JS, Jørgensen JJ, Ostrowski SR, Holtan A, Gundersen Y, Johansson PI, et al. Changes in fibrinogen availability and utilization in an animal model of traumatic coagulopathy. Scandinavian Journal of Trauma, Resuscitation and Emergency Medicine. 2013;21(1):56. doi: 10.1186/1757-7241-21-56.

129. Emery M, Nanchen N, Preitner F, Ibberson M, Roduit R. Biological characterization of gene response to insulin-induced hypoglycemia in mouse retina. PLOS ONE. 2016;11(2):e0150266. doi: 10.1371/journal.pone.0150266.

130. Holmes AM, Emmans CJ, Coleman R, Smith TE, Hosie CA. Effects of transportation, transport medium and re-housing on xenopus laevis (daudin). Gen Comp Endocrinol. 2018;266:21–8. Epub 2018/03/17. doi: 10.1016/j.ygcen.2018.03.015. PubMed PMID: 29545087.

131. Conour LA, Murray KA, Brown MJ. Preparation of animals for research--issues to consider for rodents and rabbits. ILAR J. 2006;47(4):283–93. Epub 2006/09/12. PubMed PMID: 16963809.

132. Obernier JA, Baldwin RL. Establishing an appropriate period of acclimatization following transportation of laboratory animals. ILAR Journal. 2006;47(4):364–9. doi: 10.1093/ilar.47.4.364.

133. Krahn DD, Gosnell BA, Majchrzak MJ. The anorectic effects of crh and restraint stress decrease with repeated exposures. Biol Psychiatry. 1990;27(10):1094–102. Epub 1990/05/15. PubMed PMID: 2340320.

134. Pitman DL, Ottenweller JE, Natelson BH. Plasma corticosterone levels during repeated presentation of two intensities of restraint stress: Chronic stress and habituation. Physiol Behav. 1988;43(1):47–55. Epub 1988/01/01. PubMed PMID: 3413250.

135. Brock AJ, Goody SMG, Mead AN, Sudwarts A, Parker MO, Brennan CH. Assessing the value of the zebrafish conditioned place preference model for predicting human abuse potential. J Pharmacol Exp Ther. 2017;363(1):66–79. Epub 2017/08/10. doi: 10.1124/jpet.117.242628. PubMed PMID: 28790193; PubMed Central PMCID: PMCPMC5602714.

136. Turner PV, Brabb T, Pekow C, Vasbinder MA. Administration of substances to laboratory animals: Routes of administration and factors to consider. Journal of the American Association for Laboratory Animal Science: JAALAS. 2011;50(5):600–13. PubMed PMID: PMC3189662.

137. Fueger BJ, Czernin J, Hildebrandt I, Tran C, Halpern BS, Stout D, et al. Impact of animal handling on the results of 18f-fdg pet studies in mice. J Nucl Med. 2006;47(6):999–1006. Epub 2006/06/03. PubMed PMID: 16741310.

138. Aslan Y, Tadjuidje E, Zorn AM, Cha SW. High-efficiency non-mosaic crispr-mediated knock-in and indel mutation in f0 xenopus. Development. 2017;144(15):2852–8. Epub 2017/07/12. doi: 10.1242/dev.152967. PubMed PMID: 28694259; PubMed Central PMCID: PMCPMC5560047.

139. Elmorsy S, Soliman G, Rashed L, Elgendy H. The response to sedative doses of propofol and dexmedetomidine in a prenatal valproate autistic rat model. Kasr Al Ainy Medical Journal. 2018;24(1):32–9. doi: 10.4103/kamj.kamj_37_17.

140. Ash JA, Lu H, Taxier LR, Long JM, Yang Y, Stein EA, et al. Functional connectivity with the retrosplenial cortex predicts cognitive aging in rats. Proc Natl Acad Sci U S A. 2016;113(43):12286–91. Epub 2016/10/30. doi: 10.1073/pnas.1525309113. PubMed PMID: 27791017; PubMed Central PMCID: PMCPMC5087009.

141. Altman DG. Why we need confidence intervals. World J Surg. 2005;29(5):554–6. Epub 2005/04/14. doi: 10.1007/s00268-005-7911-0. PubMed PMID: 15827844.

142. Moher D, Hopewell S, Schulz KF, Montori V, Gøtzsche PC, Devereaux PJ, et al. Consort 2010 explanation and elaboration: Updated guidelines for reporting parallel group randomised trials. BMJ. 2010;340. doi: 10.1136/bmj.c869.

143. Nakagawa S, Cuthill IC. Effect size, confidence interval and statistical significance: A practical guide for biologists. Biol Rev Camb Philos Soc. 2007;82(4):591–605. Epub 2007/10/20. doi: 10.1111/j.1469-185X.2007.00027.x. PubMed PMID: 17944619.

144. Haynes RB, Mulrow CD, Huth EJ, Altman DG, Gardner MJ. More informative abstracts revisited. Ann Intern Med. 1990;113(1):69–76. Epub 1990/07/01. PubMed PMID: 2190518.

145. Berwanger O, Ribeiro RA, Finkelsztejn A, Watanabe M, Suzumura EA, Duncan BB, et al. The quality of reporting of trial abstracts is suboptimal: Survey of major general medical journals. J Clin Epidemiol. 2009;62(4):387–92. Epub 2008/11/18. doi: 10.1016/j.jclinepi.2008.05.013. PubMed PMID: 19010643.

146. Can OS, Yilmaz AA, Hasdogan M, Alkaya F, Turhan SC, Can MF, et al. Has the quality of abstracts for randomised controlled trials improved since the release of consolidated standards of reporting trial guideline for abstract reporting? A survey of four high-profile anaesthesia journals. Eur J Anaesthesiol. 2011;28(7):485–92. Epub 2010/11/03. doi: 10.1097/EJA.0b013e32833fb96f. PubMed PMID: 21037480.

147. Pitkin RM, Branagan MA, Burmeister LF. Accuracy of data in abstracts of published research articles. JAMA. 1999;281(12):1110–1. Epub 1999/04/03. PubMed PMID: 10188662.

148. Boutron I, Altman DG, Hopewell S, Vera-Badillo F, Tannock I, Ravaud P. Impact of spin in the abstracts of articles reporting results of randomized controlled trials in the field of cancer: The spiin randomized controlled trial. J Clin Oncol. 2014;32(36):4120–6. Epub 2014/11/19. doi: 10.1200/JCO.2014.56.7503. PubMed PMID: 25403215.

149. Toledo AC, Sakoda CPP, Perini A, Pinheiro NM, Magalhães RM, Grecco S, et al. Flavonone treatment reverses airway inflammation and remodelling in an asthma murine model. British Journal of Pharmacology. 2013;168(7):1736–49. doi: 10.1111/bph.12062. PubMed PMID: 23170811.

150. DeBoer SP, Garner JP, McCain RR, Lay Jr DC, Eicher SD, Marchant-Forde JN. An initial investigation into the effects of isolation and enrichment on the welfare of laboratory pigs housed in the pigturn^®^ system, assessed using tear staining, behaviour, physiology and haematology. Animal Welfare. 2015;24(1):15–27. doi: 10.7120/09627286.24.1.015.

151. Avey MT, Fenwick N, Griffin G. The use of systematic reviews and reporting guidelines to advance the implementation of the 3rs. Journal of the American Association for Laboratory Animal Science: JAALAS. 2015;54(2):153–62. PubMed PMID: PMC4382619.

152. Hooijmans CR, Ritskes-Hoitinga M. Progress in using systematic reviews of animal studies to improve translational research. PLOS Medicine. 2013;10(7):e1001482. doi: 10.1371/journal.pmed.1001482.

153. Donner DG, Elliott GE, Beck BR, Bulmer AC, Du Toit EF. Impact of diet-induced obesity and testosterone deficiency on the cardiovascular system: A novel rodent model representative of males with testosterone-deficient metabolic syndrome (tdmets). PLOS ONE. 2015;10(9):e0138019. doi: 10.1371/journal.pone.0138019.

154. Willner P. Validation criteria for animal models of human mental disorders: Learned helplessness as a paradigm case. Progress in Neuro-Psychopharmacology and Biological Psychiatry. 1986;10(6):677–90. doi: 10.1016/0278-5846(86)90051-5.

155. Hunt RF, Girskis KM, Rubenstein JL, Alvarez–Buylla A, Baraban SC. Gaba progenitors grafted into the adult epileptic brain control seizures and abnormal behavior. Nature neuroscience. 2013;16(6):692–7. doi: 10.1038/nn.3392. PubMed PMID: PMC3665733.

156. Hamilton N, Sabroe I, Renshaw SA. A method for transplantation of human hscs into zebrafish, to replace humanised murine transplantation models. F1000Res. 2018;7:594. Epub 2019/01/05. doi: 10.12688/f1000research.14507.2. PubMed PMID: 29946444; PubMed Central PMCID: PMCPMC6008850.2.

157. Kimmelman J, Mogil JS, Dirnagl U. Distinguishing between exploratory and confirmatory preclinical research will improve translation. PLoS Biol. 2014;12(5):e1001863. doi: 10.1371/journal.pbio.1001863.

158. Camp DM, Loeffler DA, Farrah DM, Borneman JN, LeWitt PA. Cellular immune response to intrastriatally implanted allogeneic bone marrow stromal cells in a rat model of parkinson’s disease. J Neuroinflammation. 2009;6:17. Epub 2009/06/09. doi: 10.1186/1742-2094-6-17. PubMed PMID: 19500379; PubMed Central PMCID: PMCPMC2700085.

159. Falk H, Forde PF, Bay ML, Mangalanathan UM, Hojman P, Soden DM, et al. Calcium electroporation induces tumor eradication, long-lasting immunity and cytokine responses in the ct26 colon cancer mouse model. OncoImmunology. 2017;6(5):e1301332. doi: 10.1080/2162402X.2017.1301332.

160. Bayne K, Turner PV. Animal welfare standards and international collaborations. ILAR J. 2019. Epub 2019/01/10. doi: 10.1093/ilar/ily024. PubMed PMID: 30624646.

161. Redfern WS, Tse K, Grant C, Keerie A, Simpson DJ, Pedersen JC, et al. Automated recording of home cage activity and temperature of individual rats housed in social groups: The rodent big brother project. PLoS ONE. 2017;12(9):e0181068. doi: 10.1371/journal.pone.0181068.

162. Wang X, Xue Q, Yan F, Liu J, Li S, Hu S. Ulinastatin protects against acute kidney injury in infant piglets model undergoing surgery on hypothermic low-flow cardiopulmonary bypass. PLOS ONE. 2015;10(12):e0144516. doi: 10.1371/journal.pone.0144516.

163. Berghof TV, van der Klein SA, Arts JA, Parmentier HK, van der Poel JJ, Bovenhuis H. Genetic and non-genetic inheritance of natural antibodies binding keyhole limpet hemocyanin in a purebred layer chicken line. PLoS One. 2015;10(6):e0131088. Epub 2015/06/27. doi: 10.1371/journal.pone.0131088. PubMed PMID: 26114750; PubMed Central PMCID: PMCPMC4482680.

164. Bailoo JD, Murphy E, Varholick JA, Novak J, Palme R, Würbel H. Evaluation of the effects of space allowance on measures of animal welfare in laboratory mice. Scientific Reports. 2018;8(1):713. doi: 10.1038/s41598-017-18493-6.

165. Kallnik M, Elvert R, Ehrhardt N, Kissling D, Mahabir E, Welzl G, et al. Impact of ivc housing on emotionality and fear learning in male c3heb/fej and c57bl/6j mice. Mamm Genome. 2007;18(3):173–86. Epub 2007/04/14. doi: 10.1007/s00335-007-9002-z. PubMed PMID: 17431719.

166. Holmes AM, Emmans CJ, Jones N, Coleman R, Smith TE, Hosie CA. Impact of tank background on the welfare of the african clawed frog, xenopus laevis (daudin). Applied Animal Behaviour Science. 2016;185:131–6. doi: 10.1016/j.applanim.2016.09.005. PubMed PMID: WOS:000390504700018.

167. Haseman JK, Ney E, Nyska A, Rao GN. Effect of diet and animal care/housing protocols on body weight, survival, tumor incidences, and nephropathy severity of f344 rats in chronic studies. Toxicol Pathol. 2003;31(6):674–81. Epub 2003/10/31. doi: 10.1080/01926230390241927. PubMed PMID: 14585736.

168. Morris JK, Bomhoff GL, Stanford JA, Geiger PC. Neurodegeneration in an animal model of parkinson’s disease is exacerbated by a high-fat diet. American Journal of Physiology-Regulatory, Integrative and Comparative Physiology. 2010;299(4):R1082–R90. doi: 10.1152/ajpregu.00449.2010.

169. Bayol SA, Farrington SJ, Stickland NC. A maternal ‘junk food’ diet in pregnancy and lactation promotes an exacerbated taste for ‘junk food’ and a greater propensity for obesity in rat offspring. Br J Nutr. 2007;98(4):843–51. Epub 2007/08/19. doi: 10.1017/S0007114507812037. PubMed PMID: 17697422.

170. Gaskill BN, Garner JP. Stressed out: Providing laboratory animals with behavioral control to reduce the physiological effects of stress. Lab Animal. 2017;46:142. doi: 10.1038/laban.1218.

171. Robinson I, Dowdall T, Meert TF. Development of neuropathic pain is affected by bedding texture in two models of peripheral nerve injury in rats. Neurosci Lett. 2004;368(1):107–11. Epub 2004/09/03. doi: 10.1016/j.neulet.2004.06.078. PubMed PMID: 15342144.

172. Kokolus KM, Capitano ML, Lee CT, Eng JW, Waight JD, Hylander BL, et al. Baseline tumor growth and immune control in laboratory mice are significantly influenced by subthermoneutral housing temperature. Proc Natl Acad Sci U S A. 2013;110(50):20176–81. Epub 2013/11/20. doi: 10.1073/pnas.1304291110. PubMed PMID: 24248371; PubMed Central PMCID: PMCPMC3864348.

173. Duke JL, Zammit TG, Lawson DM. The effects of routine cage-changing on cardiovascular and behavioral parameters in male sprague-dawley rats. Contemp Top Lab Anim Sci. 2001;40(1):17–20. Epub 2001/04/13. PubMed PMID: 11300670.

174. Prager E, Bergstrom H, Grunberg N, Johnson L. The importance of reporting housing and husbandry in rat research. Front Behav Neurosci. 2011;5(38). doi: 10.3389/fnbeh.2011.00038.

175. Rosenbaum MD, VandeWoude S, Johnson TE. Effects of cage-change frequency and bedding volume on mice and their microenvironment. Journal of the American Association for Laboratory Animal Science: JAALAS. 2009;48(6):763–73. PubMed PMID: PMC2786931.

176. Kappel S, Hawkins P, Mendl MT. To group or not to group? Good practice for housing male laboratory mice. Animals: an Open Access Journal from MDPI. 2017;7(12):88. doi: 10.3390/ani7120088. PubMed PMID: PMC5742782.

177. Van Loo PLP, Mol JA, Koolhaas JM, Van Zutphen BFM, Baumans V. Modulation of aggression in male mice: Influence of group size and cage size. Physiology & Behavior. 2001;72(5):675–83. doi: 10.1016/S0031-9384(01)00425-5.

178. Bleich A, Hansen AK. Time to include the gut microbiota in the hygienic standardisation of laboratory rodents. Comp Immunol Microbiol Infect Dis. 2012;35(2):81–92. Epub 2012/01/20. doi: 10.1016/j.cimid.2011.12.006. PubMed PMID: 22257867.

179. Dauchy RT, Dupepe LM, Ooms TG, Dauchy EM, Hill CR, Mao L, et al. Eliminating animal facility light-at-night contamination and its effect on circadian regulation of rodent physiology, tumor growth, and metabolism: A challenge in the relocation of a cancer research laboratory. J Am Assoc Lab Anim Sci. 2011;50(3):326–36. Epub 2011/06/07. PubMed PMID: 21640027; PubMed Central PMCID: PMCPMC3103282.

180. Chapillon P, Manneché C, Belzung C, Caston J. Rearing environmental enrichment in two inbred strains of mice: 1. Effects on emotional reactivity. Behavior Genetics. 1999;29(1):41–6. doi: 10.1023/a:1021437905913.

181. Hendershott TR, Cronin ME, Langella S, McGuinness PS, Basu AC. Effects of environmental enrichment on anxiety-like behavior, sociability, sensory gating, and spatial learning in male and female c57bl/6j mice. Behavioural Brain Research. 2016;314:215–25. doi: 10.1016/j.bbr.2016.08.004.

182. Garner JP. Stereotypies and other abnormal repetitive behaviors: Potential impact on validity, reliability, and replicability of scientific outcomes. Ilar j. 2005;46(2):106–17. Epub 2005/03/19. PubMed PMID: 15775020.

183. Gross AN-M, Engel AKJ, Würbel H. Simply a nest? Effects of different enrichments on stereotypic and anxiety-related behaviour in mice. Applied Animal Behaviour Science. 2011;134(3):239–45. doi: 10.1016/j.applanim.2011.06.020.

184. Wurbel H. Ideal homes? Housing effects on rodent brain and behaviour. Trends Neurosci. 2001;24(4):207–11. Epub 2001/03/16. PubMed PMID: 11250003.

185. Auvergne R, Déan C, El Bahh B, Arthaud S, lespinet-najib v, Rougier A, et al. Delayed kindling epileptogenesis and increased neurogenesis in adult rats housed in an enriched environment 2002. 277–85 p.

186. Salvarrey-Strati A, Watson L, Blanchet T, Lu N, Glasson SS. The influence of enrichment devices on development of osteoarthritis in a surgically induced murine model. Ilar j. 2008;49(4):23–30. Epub 2008/10/14. PubMed PMID: 18849588.

187. Hannan AJ. Environmental enrichment and brain repair: Harnessing the therapeutic effects of cognitive stimulation and physical activity to enhance experience-dependent plasticity. Neuropathol Appl Neurobiol. 2014;40(1):13–25. Epub 2013/12/21. doi: 10.1111/nan.12102. PubMed PMID: 24354721.

188. Sorge RE, Martin LJ, Isbester KA, Sotocinal SG, Rosen S, Tuttle AH, et al. Olfactory exposure to males, including men, causes stress and related analgesia in rodents. Nature Methods. 2014;11:629. doi: 10.1038/nmeth.2935

189. Kotloski RJ, Sutula TP. Environmental enrichment: Evidence for an unexpected therapeutic influence. Exp Neurol. 2015;264:121–6. Epub 2014/12/09. doi: 10.1016/j.expneurol.2014.11.012. PubMed PMID: 25483395.

190. Jensen TL, Kiersgaard MK, Sorensen DB, Mikkelsen LF. Fasting of mice: A review. Lab Anim. 2013;47(4):225–40. Epub 2013/09/13. doi: 10.1177/0023677213501659. PubMed PMID: 24025567.

191. National Research Council. Guide for the care and use of laboratory animals. 8th ed: The National Academies Press; 2011.

192. European Commission. Directive 2010/63/eu of the european parliament and of the council of 22 september 2010 on the protection of animals used for scientific purposes. Official Journal of the European Union [Internet]. 2010; 53. Available from: https://eur-lex.europa.eu/legal-content/EN/TXT/?uri=celex%3A32010L0063.

193. Heykants M, Mahabir E. Estrous cycle staging before mating led to increased efficiency in the production of pseudopregnant recipients without negatively affecting embryo transfer in mice. Theriogenology. 2016;85(5):813–21. Epub 2015/11/29. doi: 10.1016/j.theriogenology.2015.10.027. PubMed PMID: 26613855.

194. Gallastegui A, Cheung J, Southard T, Hume KR. Volumetric and linear measurements of lung tumor burden from non-gated micro-ct imaging correlate with histological analysis in a genetically engineered mouse model of non-small cell lung cancer. Lab Anim. 2018:23677218756457. Epub 2018/02/14. doi: 10.1177/0023677218756457. PubMed PMID: 29436921.

195. Jirkof P. Side effects of pain and analgesia in animal experimentation. Lab Anim (NY). 2017;46(4):123–8. Epub 2017/03/23. doi: 10.1038/laban.1216. PubMed PMID: 28328895.

196. Carbone L, Austin J. Pain and laboratory animals: Publication practices for better data reproducibility and better animal welfare. PLoS One. 2016;11(5):e0155001. Epub 2016/05/14. doi: 10.1371/journal.pone.0155001. PubMed PMID: 27171143; PubMed Central PMCID: PMCPMC4865140.

197. Gaspani L, Bianchi M, Limiroli E, Panerai AE, Sacerdote P. The analgesic drug tramadol prevents the effect of surgery on natural killer cell activity and metastatic colonization in rats. J Neuroimmunol. 2002;129(1-2):18–24. Epub 2002/08/06. PubMed PMID: 12161016.

198. Getty CM, Dilger RN. Moderate perinatal choline deficiency elicits altered physiology and metabolomic profiles in the piglet. PLoS One. 2015;10(7):e0133500. Epub 2015/07/22. doi: 10.1371/journal.pone.0133500. PubMed PMID: 26196148; PubMed Central PMCID: PMCPMC4510435.

199. Teilmann AC, Falkenberg MK, Hau J, Abelson KS. Comparison of silicone and polyurethane catheters for the catheterization of small vessels in mice. Lab Anim (NY). 2014;43(11):397–403. Epub 2014/10/22. doi: 10.1038/laban.570. PubMed PMID: 25333592.

200. Bauer G, Dao MA, Case SS, Meyerrose T, Wirthlin L, Zhou P, et al. In vivo biosafety model to assess the risk of adverse events from retroviral and lentiviral vectors. Mol Ther. 2008;16(7):1308–15. Epub 2008/05/08. doi: 10.1038/mt.2008.93. PubMed PMID: 18461052; PubMed Central PMCID: PMCPMC3013368.

201. Bellanger S, Benrezzak O, Battista MC, Naimi F, Labbe SM, Frisch F, et al. Experimental dog model for assessment of fasting and postprandial fatty acid metabolism: Pitfalls and feasibility. Lab Anim. 2015;49(3):228–40. Epub 2015/01/08. doi: 10.1177/0023677214566021. PubMed PMID: 25563731.

202. Hendriksen C, Morton D, Cussler K. Use of humane endpoints to minimise suffering. Howard B, T N, Peratta G, editors: CRC Press, Florida, USA; 2011.

203. Percie du Sert N, Alfieri A, Allan SM, Carswell HV, Deuchar GA, Farr TD, et al. The improve guidelines (ischaemia models: Procedural refinements of in vivo experiments). J Cereb Blood Flow Metab. 2017;37(11):3488–517. Epub 2017/08/12. doi: 10.1177/0271678X17709185. PubMed PMID: 28797196; PubMed Central PMCID: PMCPMC5669349.

204. Morton DB. A systematic approach for establishing humane endpoints. ILAR Journal. 2000;41(2):80–6. doi: 10.1093/ilar.41.2.80.

205. Muscella A, Vetrugno C, Cossa LG, Antonaci G, De Nuccio F, De Pascali SA, et al. In vitro and in vivo antitumor activity of [pt(o,o’-acac)(gamma-acac)(dms)] in malignant pleural mesothelioma. PLoS One. 2016;11(11):e0165154. Epub 2016/11/03. doi: 10.1371/journal.pone.0165154. PubMed PMID: 27806086; PubMed Central PMCID: PMCPMC5091852.

206. Glasziou P, Altman DG, Bossuyt P, Boutron I, Clarke M, Julious S, et al. Reducing waste from incomplete or unusable reports of biomedical research. Lancet. 2014;383(9913):267–76. Epub 2014/01/15. doi: 10.1016/s0140-6736(13)62228-x. PubMed PMID: 24411647.

207. Schirmer B, Wedekind D, Glage S, Neumann D. Deletion of il-18 expression ameliorates spontaneous kidney failure in mrllpr mice. PLOS ONE. 2015;10(10):e0140173. doi: 10.1371/journal.pone.0140173.

208. Ameen-Ali KE, Eacott MJ, Easton A. A new behavioural apparatus to reduce animal numbers in multiple types of spontaneous object recognition paradigms in rats. Journal of neuroscience methods. 2012;211(1):66–76. doi: 10.1016/j.jneumeth.2012.08.006. PubMed PMID: 22917958.

209. Banwell V, Sena ES, Macleod MR. Systematic review and stratified meta-analysis of the efficacy of interleukin-1 receptor antagonist in animal models of stroke. Journal of Stroke and Cerebrovascular Diseases. 2009;18(4):269–76. doi: 10.1016/j.jstrokecerebrovasdis.2008.11.009.

210. Pradillo JM, Denes A, Greenhalgh AD, Boutin H, Drake C, McColl BW, et al. Delayed administration of interleukin-1 receptor antagonist reduces ischemic brain damage and inflammation in comorbid rats. Journal of Cerebral Blood Flow & Metabolism. 2012;32(9):1810–9. doi: 10.1038/jcbfm.2012.101.

211. Ioannidis JP. Limitations are not properly acknowledged in the scientific literature. J Clin Epidemiol. 2007;60(4):324–9. Epub 2007/03/10. doi: 10.1016/j.jclinepi.2006.09.011. PubMed PMID: 17346604.

212. Wieschowski S, Chin WWL, Federico C, Sievers S, Kimmelman J, Strech D. Preclinical efficacy studies in investigator brochures: Do they enable risk–benefit assessment? PLOS Biology. 2018;16(4):e2004879. doi: 10.1371/journal.pbio.2004879.

213. Diaz A, Perez A, Sreevatsan S, Davies P, Culhane M, Torremorell M. Association between influenza a virus infection and pigs subpopulations in endemically infected breeding herds. PLOS ONE. 2015;10(6):e0129213. doi: 10.1371/journal.pone.0129213.

214. Dhondup Y, Sjaastad I, Scott H, Sandanger Ø, Zhang L, Haugstad SB, et al. Sustained toll-like receptor 9 activation promotes systemic and cardiac inflammation, and aggravates diastolic heart failure in serca2a ko mice. PLOS ONE. 2015;10(10):e0139715. doi: 10.1371/journal.pone.0139715.

215. Chalmers I, Bracken MB, Djulbegovic B, Garattini S, Grant J, Gülmezoglu AM, et al. How to increase value and reduce waste when research priorities are set. The Lancet. 2014;383(9912):156–65. doi: 10.1016/S0140-6736(13)62229-1.

216. Voelkl B, Vogt L, Sena ES, Würbel H. Reproducibility of preclinical animal research improves with heterogeneity of study samples. PLOS Biology. 2018;16(2):e2003693. doi: 10.1371/journal.pbio.2003693.

217. Munafò MR, Davey Smith G. Robust research needs many lines of evidence. Nature. 2018;553(7689):399–401. Epub 2018/01/26. doi: 10.1038/d41586-018-01023-3. PubMed PMID: 29368721.

218. Van Den Berge N, Vanhove C, Descamps B, Dauwe I, van Mierlo P, Vonck K, et al. Functional mri during hippocampal deep brain stimulation in the healthy rat brain. PLOS ONE. 2015;10(7):e0133245. doi: 10.1371/journal.pone.0133245.

219. Sacilotto N, Castillo J, Riffo-Campos ÁL, Flores JM, Hibbitt O, Wade-Martins R, et al. Growth arrest specific 1 (gas1) gene overexpression in liver reduces the in vivo progression of murine hepatocellular carcinoma and partially restores gene expression levels. PLOS ONE. 2015;10(7):e0132477. doi: 10.1371/journal.pone.0132477.

220. Chambers CD, Feredoes E, Muthukumaraswamy SD, Etchells PJ. Instead of “playing the game” it is time to change the rules: Registered reports at *aims neuroscience* and beyond. AIMS Neuroscience. 2014;1(1):4–17. doi: DOI: 10.3934/Neuroscience2014.1.4.

221. Nosek BA, Lakens D. Registered reports a method to increase the credibility of published results. Social Psychology. 2014;45(3):137–41. doi: 10.1027/1864-9335/a000192. PubMed PMID: WOS:000336836900001.

222. Kaplan RM, Irvin VL. Likelihood of null effects of large nhlbi clinical trials has increased over time. Plos One. 2015;10(8). doi: ARTN e0132382 10.1371/journal.pone.0132382. PubMed PMID: WOS:000359061400022.

223. Allen C, Mehler D. Open science challenges, benefits and tips in early career and beyond. PsyArXiv. 2018. doi: 10.31234/osf.io/3czyt.

224. Nosek BA, Ebersole CR, DeHaven AC, Mellor DT. The preregistration revolution. Proc Natl Acad Sci U S A. 2018;115(11):2600–6. Epub 2018/03/14. doi: 10.1073/pnas.1708274114. PubMed PMID: 29531091; PubMed Central PMCID: PMCPMC5856500.

225. Macleod M, Howells D. Protocols for laboratory research. Evidence-based Preclinical Medicine. 2016;3(2):e00021. doi: doi:10.1002/ebm2.21.

226. Mantis C, Kandela I, Aird F, Reproducibility Project: Cancer B. Replication study: Coadministration of a tumor-penetrating peptide enhances the efficacy of cancer drugs. eLife. 2017;6:e17584. doi: 10.7554/eLife.17584.

227. Jeronimo S, Khadraoui M, Wang DP, Martin K, Lesku JA, Robert KA, et al. Plumage color manipulation has no effect on social dominance or fitness in zebra finches. Behavioral Ecology. 2018;29(2):459–67. doi: 10.1093/beheco/arx195. PubMed PMID: WOS:000427885600027.

228. Sheldon EL, Griffith SC. Embryonic heart rate predicts prenatal development rate, but is not related to post-natal growth rate or activity level in the zebra finch (taeniopygia guttata). Ethology. 2018;124(11):829–37. doi: 10.1111/eth.12817. PubMed PMID: WOS:000449485800006.

229. Kafkafi N, Mayo CL, Elmer GI. Mining mouse behavior for patterns predicting psychiatric drug classification. Psychopharmacology (Berl). 2014;231(1):231–42. Epub 2013/08/21. doi: 10.1007/s00213-013-3230-6. PubMed PMID: 23958942.

230. Stodden V, Guo P, Ma Z. Toward reproducible computational research: An empirical analysis of data and code policy adoption by journals. PLOS ONE. 2013;8(6):e67111. doi: 10.1371/journal.pone.0067111.

231. Piwowar HA, Day RS, Fridsma DB. Sharing detailed research data is associated with increased citation rate. PLoS ONE. 2007;2(3):e308. doi: 10.1371/journal.pone.0000308.

232. DataCitationSynthesis Group. Joint declaration of data citation principles San Diego CA: FORCE11; 2014 [cited 2018 22 May].

233. Wilkinson MD, Dumontier M, Aalbersberg IJ, Appleton G, Axton M, Baak A, et al. The fair guiding principles for scientific data management and stewardship. Scientific Data. 2016;3:160018. doi: 10.1038/sdata.2016.18.

234. Federer LM, Lu Y-L, Joubert DJ, Welsh J, Brandys B. Biomedical data sharing and reuse: Attitudes and practices of clinical and scientific research staff. PLoS ONE. 2015;10(6):e0129506. doi: 10.1371/journal.pone.0129506.

235. Mamrot J, Legaie R, Ellery SJ, Wilson T, Seemann T, Powell DR, et al. De novo transcriptome assembly for the spiny mouse (acomys cahirinus). Scientific Reports. 2017;7(1):8996. doi: 10.1038/s41598-017-09334-7.

236. Bero L, Anglemyer A, Vesterinen H, Krauth D. The relationship between study sponsorship, risks of bias, and research outcomes in atrazine exposure studies conducted in non-human animals: Systematic review and meta-analysis. Environment International. 2016;92-93:597–604. doi: 10.1016/j.envint.2015.10.011.

237. Dalbøge LS, Pedersen PJ, Hansen G, Fabricius K, Hansen HB, Jelsing J, et al. A hamster model of diet-induced obesity for preclinical evaluation of anti-obesity, anti-diabetic and lipid modulating agents. PLOS ONE. 2015;10(8):e0135634. doi: 10.1371/journal.pone.0135634.

238. Garcia de la serrana D, Vieira VLA, Andree KB, Darias M, Estévez A, Gisbert E, et al. Development temperature has persistent effects on muscle growth responses in gilthead sea bream. PLOS ONE. 2012;7(12):e51884. doi: 10.1371/journal.pone.0051884.

239. Lundh A, Sismondo S, Lexchin J, Busuioc OA, Bero L. Industry sponsorship and research outcome. Cochrane Database of Systematic Reviews. 2012;(12). doi: 10.1002/14651858.MR000033.pub2. PubMed PMID: MR000033.

240. Popelut A, Valet F, Fromentin O, Thomas A, Bouchard P. Relationship between sponsorship and failure rate of dental implants: A systematic approach. PLOS ONE. 2010;5(4):e10274. doi: 10.1371/journal.pone.0010274.

241. Lexchin J, Bero LA, Djulbegovic B, Clark O. Pharmaceutical industry sponsorship and research outcome and quality: Systematic review. Bmj. 2003;326(7400):1167–70. Epub 2003/05/31. doi: 10.1136/bmj.326.7400.1167. PubMed PMID: 12775614; PubMed Central PMCID: PMCPMC156458.

242. Krauth D, Anglemyer A, Philipps R, Bero L. Nonindustry-sponsored preclinical studies on statins yield greater efficacy estimates than industry-sponsored studies: A meta-analysis. PLoS Biology. 2014;12(1):e1001770. doi: 10.1371/journal.pbio.1001770.

243. Genchi M, Prati P, Vicari N, Manfredini A, Sacchi L, Clementi E, et al. Francisella tularensis: No evidence for transovarial transmission in the tularemia tick vectors dermacentor reticulatus and ixodes ricinus. PLOS ONE. 2015;10(8):e0133593. doi: 10.1371/journal.pone.0133593.

244. Kolahian S, Sadri H, Shahbazfar AA, Amani M, Mazadeh A, Mirani M. The effects of leucine, zinc, and chromium supplements on inflammatory events of the respiratory system in type 2 diabetic rats. PLOS ONE. 2015;10(7):e0133374. doi: 10.1371/journal.pone.0133374.

245. Eyre-Walker A, Stoletzki N. The assessment of science: The relative merits of post-publication review, the impact factor, and the number of citations. PLOS Biology. 2013;11(10):e1001675. doi: 10.1371/journal.pbio.1001675.

